# Disome-seq reveals widespread ribosome collisions that recruit co-translational chaperones

**DOI:** 10.1101/746875

**Authors:** Taolan Zhao, Yan-Ming Chen, Yu Li, Jia Wang, Siyu Chen, Ning Gao, Wenfeng Qian

**Affiliations:** State Key Laboratory of Plant Genomics, Institute of Genetics and Developmental Biology, Chinese Academy of Sciences, Beijing 100101, China; Key Laboratory of Genetic Network Biology, Institute of Genetics and Developmental Biology, Chinese Academy of Sciences, Beijing 100101, China; University of Chinese Academy of Sciences, Beijing 100049, China; Peking University-Tsinghua University-National Institute of Biological Sciences Joint Graduate Program, School of Life Science, Tsinghua University, Beijing 100084, China; State Key Laboratory of Membrane Biology, Peking-Tsinghua Center for Life Sciences, School of Life Sciences, Peking University, Beijing 100871, China

**Keywords:** Translation elongation, disome-seq, ribosome collisions, translational pausing, ribosome release, disome structure, ribosome-associated chaperones, co-translational protein folding, protein homeostasis

## Abstract

Regulation of translation elongation plays a crucial role in determining absolute protein levels and ensuring the correct localization and folding of proteins. Much of our knowledge regarding translation elongation comes from the sequencing of mRNA fragments protected by single ribosomes (ribo-seq). However, larger protected mRNA fragments have been observed, suggesting the existence of an alternative and previously hidden layer of regulation. In this study, we performed disome-seq to sequence mRNA fragments protected by two stacked ribosomes — a product of translational pauses during which the 5′-ribosome collides with the 3′-paused one. We detected widespread ribosome collisions that are missed in traditional ribo-seq. These collisions are due to 1) slow ribosome release when stop codons are at the A-site, 2) slow peptide bond formation from proline, glycine, asparagine, and cysteine when they are at the P-site, and 3) slow leaving of polylysine from the exit tunnel of ribosomes. The paused ribosomes can continue translating after collisions, as suggested by the structure of disomes obtained by cryo-electron microscopy (cryo-EM). Collided ribosomes recruit chaperones, which can aid in the co-translational folding of the nascent peptides. Therefore, cells use regulated ribosome collisions to ensure protein homeostasis.

## INTRODUCTION

Translation elongation is a crucial process through which the genetic information in a transcript is sequentially decoded to a peptide chain by ribosomes. Yet the mRNA sequence of coding regions can harbor more information than the amino-acid sequence (Chu et al., 2014); the local rate of translation elongation is non-uniform and fine-tuned (Stein and Frydman, 2019). A change in the rate of translation elongation can lead to developmental abnormalities, neurologic diseases, and cancers (Richter and Coller, 2015).

Despite the importance of translation elongation, it has been mainly studied with heterologous reporter genes (Ikeuchi et al., 2019; Juszkiewicz and Hegde, 2017; Sundaramoorthy et al., 2017). Strong ribosomal stalling signal was placed in these reporter genes; the 5′-ribosome collides into the stalled ribosome, leading to a di-ribosome (we hereafter refer such stacked ribosomes induced by strong ribosomal stalling signal in heterologous reporters as to di-ribosomes). Structure analyses indicated that di-ribosomes were often unable to resume translation and were disassembled through triggering the ribosome-associated protein quality control (RQC) pathway (Brandman and Hegde, 2016; Ikeuchi et al., 2019; Joazeiro, 2019; Park and Subramaniam, 2019).

The knowledge of the causes for endogenous translational pausing remains highly limited, mainly because the detection of translational pauses is technically challenging in endogenous genes. The development of ribo-seq, an approach that sequences ribosome protected mRNA fragments at codon resolution, significantly increased our knowledge on translation elongation (Ingolia, 2014; Ingolia et al., 2009). Accumulation of ribosome footprints at a site indicates slow translation elongation (i.e., a translational pause); based on this idea, sequence determinants of translation elongation have been discovered, such as synonymous codon usage (Hussmann et al., 2015; Qian et al., 2012; Weinberg et al., 2016), positively charged amino acids (Charneski and Hurst, 2013), and mRNA secondary structures (Yang et al., 2014). However, traditional ribo-seq misses the information of ribosome collisions (Guydosh and Green, 2014; Ivanov et al., 2018). Instead, ribosome collisions can be studied by sequencing the mRNA fragments protected by disomes, which refer to endogenous stacked ribosomes in this study. Disomes were detectable by sucrose gradient centrifugation (Diament et al., 2018) and were observed in faulty mRNAs or 3′-untranslated regions (Guydosh and Green, 2014). However, the genomic landscape and the sequence determinants of endogenous ribosome collisions remain largely unknown in the coding sequences of faithfully transcribed mRNAs.

The consequence of endogenous translational pausing also remains unclear. It has been suggested that translation elongation can regulate co-translational protein folding (Sharma and O’Brien, 2018; Stein and Frydman, 2019; Wruck et al., 2017). For example, accumulating evidence supported that the CAG expansion in Huntington’s disease led to incorrect translational pausing and therefore, incorrect folding of the signal peptide for the subcellular localization of the Htt protein (Nissley and O’Brien, 2016). Non-optimal codons formed clusters during evolution (Chen et al., 2017; Clarke and Clark, 2008); they may create slow-translation regions and participate in the regulation of protein folding (Yu et al., 2015). However, the mechanisms by which translational pauses regulate co-translational protein folding remain understudied.

In this study, we captured the mRNA footprints that were protected by two stacked ribosomes in fast proliferating yeast cells. Such data provide a unique chance to reveal the translational dynamics that are undetectable by the traditional ribo-seq (i.e., monosome-seq). With bioinformatics analyses we identify the sequence features that are associated with ribosome collision. Cryo-EM analyses indicate that the translation of colliding ribosomes can be resumed. In fact, collided ribosomes often recruit chaperones to assist protein folding, as indicated by mass spectrometry analyses. As a consequence, a nascent peptide is ready to be correctly folded during translation.

## RESULTS

### Translational pauses generate disomes from collisions of ribosomes

Most mRNAs are associated with multiple ribosomes (Arava et al., 2003), and the speed of translational elongation varies, with some events such as tRNA depletion, known to cause ribosomes to slow down and even to pause translation (Richter and Coller, 2015). We therefore hypothesized that a slowdown or pause of one downstream ribosome might generate a collision between the paused ribosome and the 5′-elongating ribosome. These collisions would result in a single RNase I resistant fragment with twice the footprint length of a single ribosome **(Figure 1A)**.

**Figure 1.**
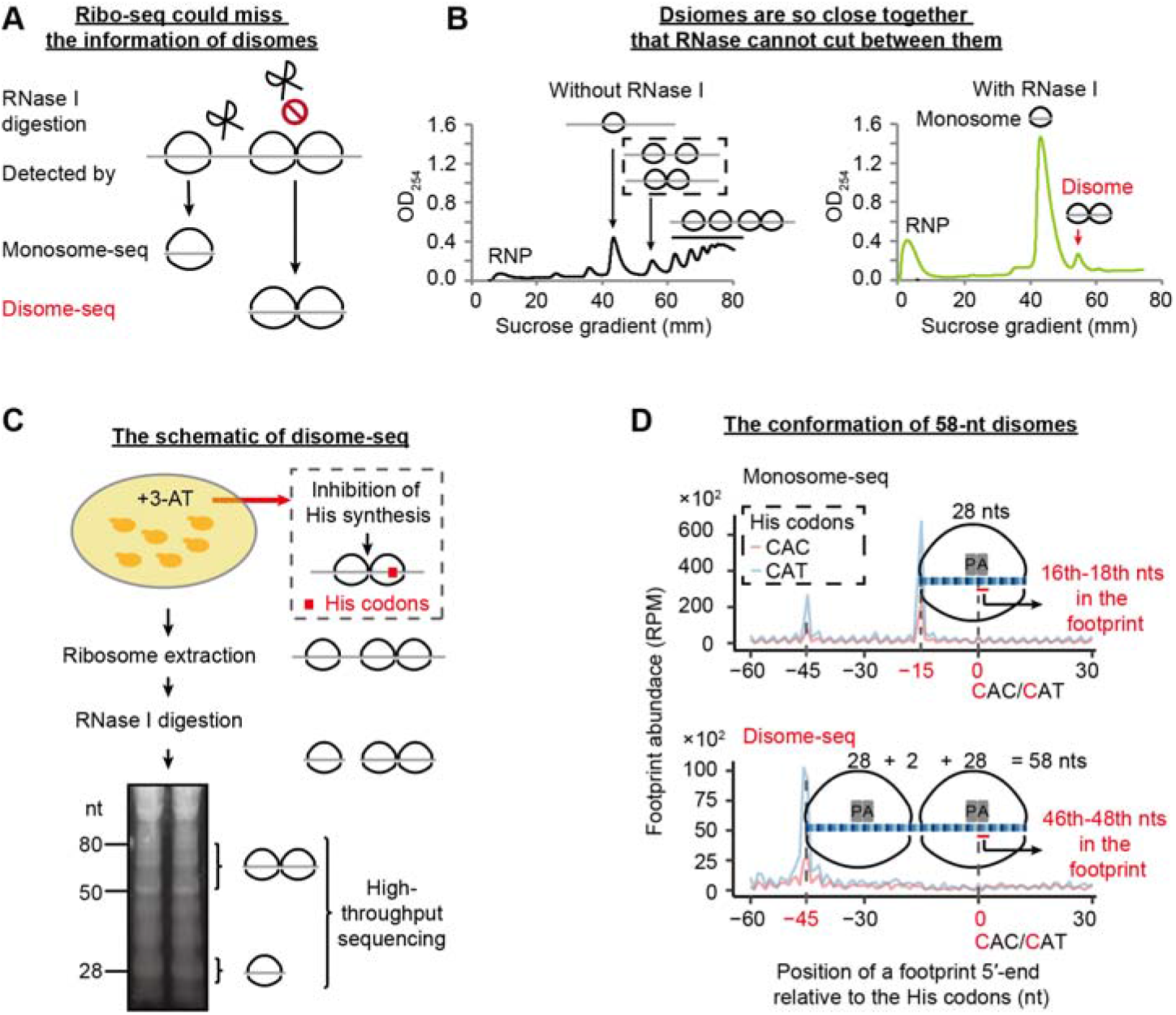
Disome-seq detects ribosome collisions. (A) A schematic graph explains why traditional ribo-seq could miss the information on ribosome collisions and why the investigation of disome footprints may provide unique information on translation. (B) The disome persisted after RNase I digestion. Sucrose gradient profiles of the ribosome-mRNA complexes without (black) and with (green) RNase I digestion are shown. The *x*-axis displays the positions in the 5-50% sucrose gradient. The *y*-axis indicates the RNA abundance inferred from UV absorption (OD_254_). RNP, free ribonucleoprotein. (C) The schematic of disome-seq for 3-AT treated yeast cells. (D) Determination of the disome conformation. Aggregated abundance profiles of the 5′-end of monosome (top) and disome footprints (bottom) are plotted around histidine (His) codons. Footprints were aligned by the first nucleotide of His codons (set at position 0). P and A represent the P-site and A-site of a ribosome, respectively. The 5′-end of the monosome (disome) footprints exhibited the main peak 15-nt (45-nt) upstream of the histidine codons in the yeast genome, indicating that the A-site of the ribosome (3′-ribosome in a disome) locates at the 16^th^–18^th^ (46^th^–48^th^) nts in the 28-nt monosome (58-nt disome) footprints. The average abundance of replicates is shown.

To test this idea, we extracted ribosome-bound mRNA from exponentially dividing yeast cells, digested the unprotected mRNA using RNase I, and performed sucrose gradient ultracentrifugation to separate particles of different densities. In addition to the abundant monosome particles, we observed a significant amount of particles whose density was the same as that of pre-digestion transcripts bound by two ribosomes, suggesting that the particle contains two ribosomes (**Figure 1B**). These disomes persisted with increased RNase I concentration, indicating that they are not the result of incomplete digestion **(Figure S1)**.

To determine if disomes were caused by paused ribosomes, we induced translational pauses at histidine codons by growing yeast in a low dose of 3-Amino-1,2,4-triazole (3-AT), an inhibitor of histidine biosynthesis (Klopotowski and Wiater, 1965) for ~ 2 generations, and performed high-throughput sequencing on monosome (monosome-seq) and disome (disome-seq) fragments (**Figure 1C** and **Figure S2A)**. Consistent with previous observations (Ingolia et al., 2009), in-frame 28-nucleotide (nt) footprints were most abundant in the monosome library (**Figure S3A** and **Table S1**). In contrast, the most abundant footprints in disome-seq were 58 and 59-nt, with the 58-nt peak being in-frame (**Figure S3B** and **Table S2**). As a negative control, mRNA-seq reads did not display any 3-nt periodicity (**Figure S3C** and **Table S3**).

During histidine starvation, ribosomes should be paused when the histidine codons CAC and CAT are at the A-site, where codon-anticodon recognition takes place. Consistently, the main peak in the 28-nt monosome footprints was 15-nt upstream of the histidine codons (**Figure 1D**). The main peak in the 58-nt disome footprints was 45-nt upstream of the histidine codons, 30-nt upstream of the 15-nt peak from the monosome footprints (**Figure 1D**). This 30-nt spacing between the two peaks perfectly fits one in-frame ribosome, suggesting that the 58-nt disome footprints were composed of two collided ribosomes of which the downstream one was paused. The sharp peak in **Figure 1D** also indicates that disome-seq detects ribosome collisions at codon resolution.

### Disome-seq enables detection of widespread translational pauses which cannot be identified via monosome sequencing

To identify the genomic locations of ribosome collisions in fast-proliferating cells, we performed monosome-seq and disome-seq in yeast cells growing in the mid-log phase in the rich medium (**Figure 2A**, **Figures S2B, S3D–I** and **Tables S1-S3**). Ribosome collisions were observed in 3929 out of the 5349 translated genes (73%, **Figure 2B**). This proportion remained substantial (24%, 1156/4742) when we applied a more stringent criterion — a ribosome collision event was called when at least two unique molecular identifiers (UMIs) existed in both biological replicates of disome-seq. These observations indicate widespread ribosome collisions in unstressed cells.

**Figure 2.**
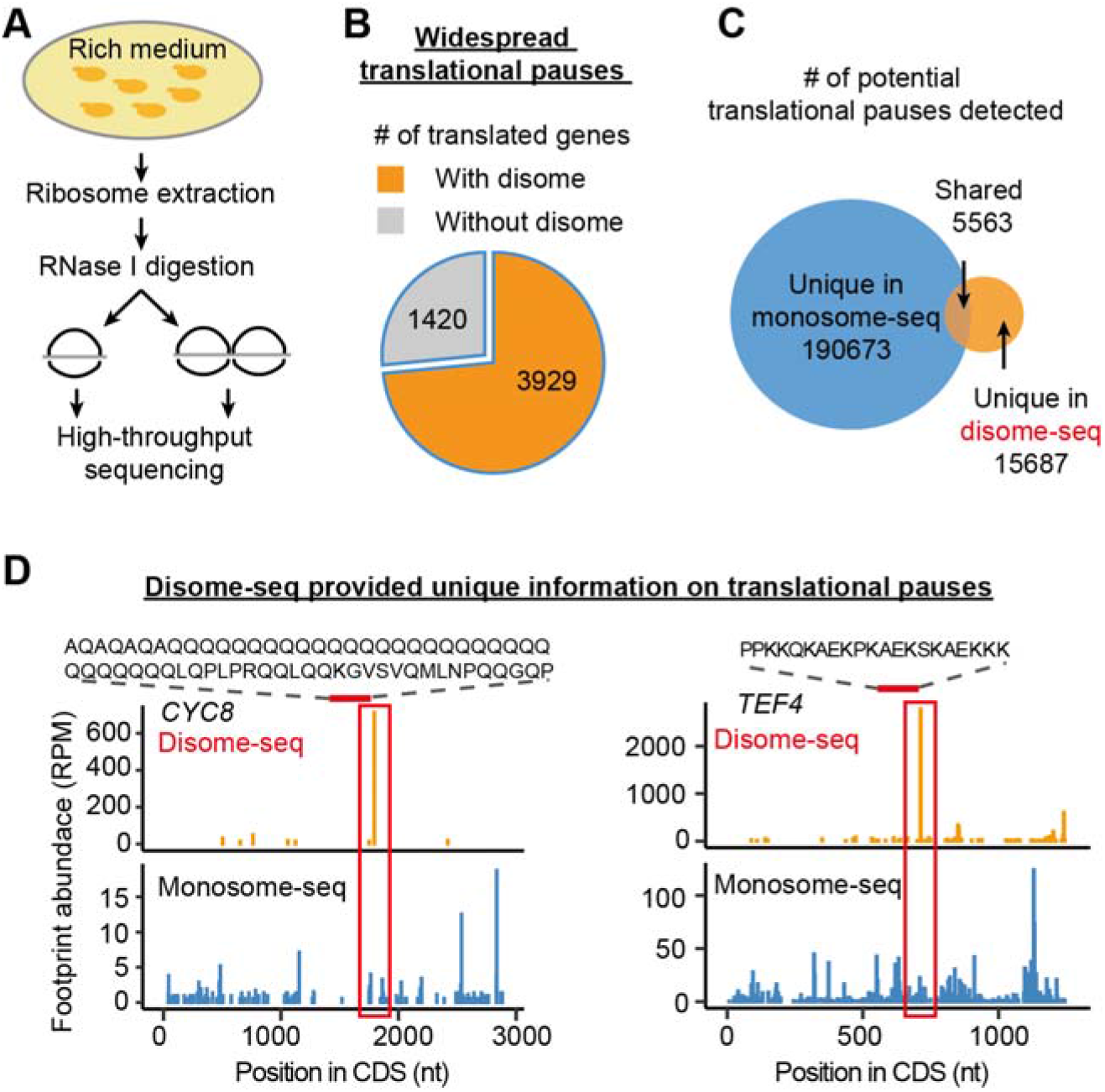
Disome-seq detects the translational pauses missed in monosome-seq. (A) The schematic of disome-seq for yeast cells cultivated in the rich medium. (B) Widespread ribosome collisions were detected in the yeast genome. Numbers of translated genes (genes with at least one monosome footprint) with and without disome footprints are shown in orange and grey, respectively. To generate this panel, the reads in biological replicates were combined. (C) The potential translational pauses detected from monosome footprints (blue) and disome footprints (orange) rarely overlapped. Only the codon site with the footprint abundance greater than the mean of the corresponding gene was considered as a potential translational pause. If the A-site of a monosome footprint overlaps the A-site of the 3′-ribosome of a disome footprint, the translational pause is “shared.” (D) Two genes exemplify the unique information of translational pauses obtained by disome-seq. The A-site of a monosome footprint (blue) or that of the 3′-ribosome of a disome footprint (orange) is shown along the coding sequence (CDS) of *CYC8* and *TEF4*.

There are two possibilities regarding ribosome collisions inside open reading frames and the relation between translational pauses identified in monosome-seq vs. disome-seq. The first is that sites with high monosome footprint abundance identify all paused ribosomes in the cell and that the collisions identified in disome-seq are simply a subset of them. In this case, it is likely that the upstream ribosome is often far 5′ of the paused ribosome, and by the time the upstream ribosome approaches, most paused ribosome has resumed elongation. In this model, all disome-seq peaks will also be identified by monosome-seq. The second possibility is that disome-seq captures ribosome collisions that occur at locations not identified by monosome sequencing, possibly because the collision often occurs not long after the pause of the downstream ribosome.

To differentiate these two possibilities, we measured the intersection of the translational pausing events from the two methods. Disome-seq often identified translational pauses that are undetectable in monosome-seq (**Figure 2C**). For example, *CYC8* is a gene encoding a general transcriptional co-repressor that can fold as the prion [OCT+] (Patel et al., 2009); a collision between ribosomes downstream of polyglutamine (polyQ) was detected by disome-seq instead of monosome-seq (**Figure 2D**). Similarly, TEF4p, the γ subunit of elongation factor eEF1B, contains a lysine-rich region that is often ubiquitinylated or succinylated (Fang et al., 2014; Weinert et al., 2013); the downstream ribosome collision was uniquely detected by disome-seq (**Figure 2D**). The inability of monosome-seq to fully characterize the dynamics of the ribosome along the mRNA is likely due to the omission of the disome protected mRNA fragments (**Figure 1A**). Consistently, while the monosome or disome density correlated well among genes between biological replicates, the correlation between the monosome and disome densities was much weaker (**Figure S2**).

### Stop codons promote ribosome collisions

Visually, we noticed that many highly expressed genes exhibited large numbers of disome-seq reads at the stop codon, but also at internal codons, suggesting that ribosome collisions occur both internally and at the ends of open reading frames **(Figure 3A)**. To identify the causes of these collisions we searched for the sequence features at the A-site, P-site, and exit tunnel of the 3′-pausing ribosome of a disome, respectively. We defined the propensity of a codon to induce ribosome collisions (i.e., the A-site pausing score) for each of the 64 codons as its enrichment in the disome footprints (**Figure 3B**). Taking the codon GAA for example (**Figure 3B**), all disome fragments on one gene were classified into two categories based on the codon identity at the A-site, GAA or the others. The odds ratio was estimated for each gene with the corresponding codon frequency in the transcript as a control; the A-site pausing score was defined as the common odds ratio across all genes calculated by the Mantel-Haenszel test (**Figure 3B**). A significantly > 1 A-site pausing score (*P*-value < 0.01) implies a slow recognition of the codon.

**Figure 3.**
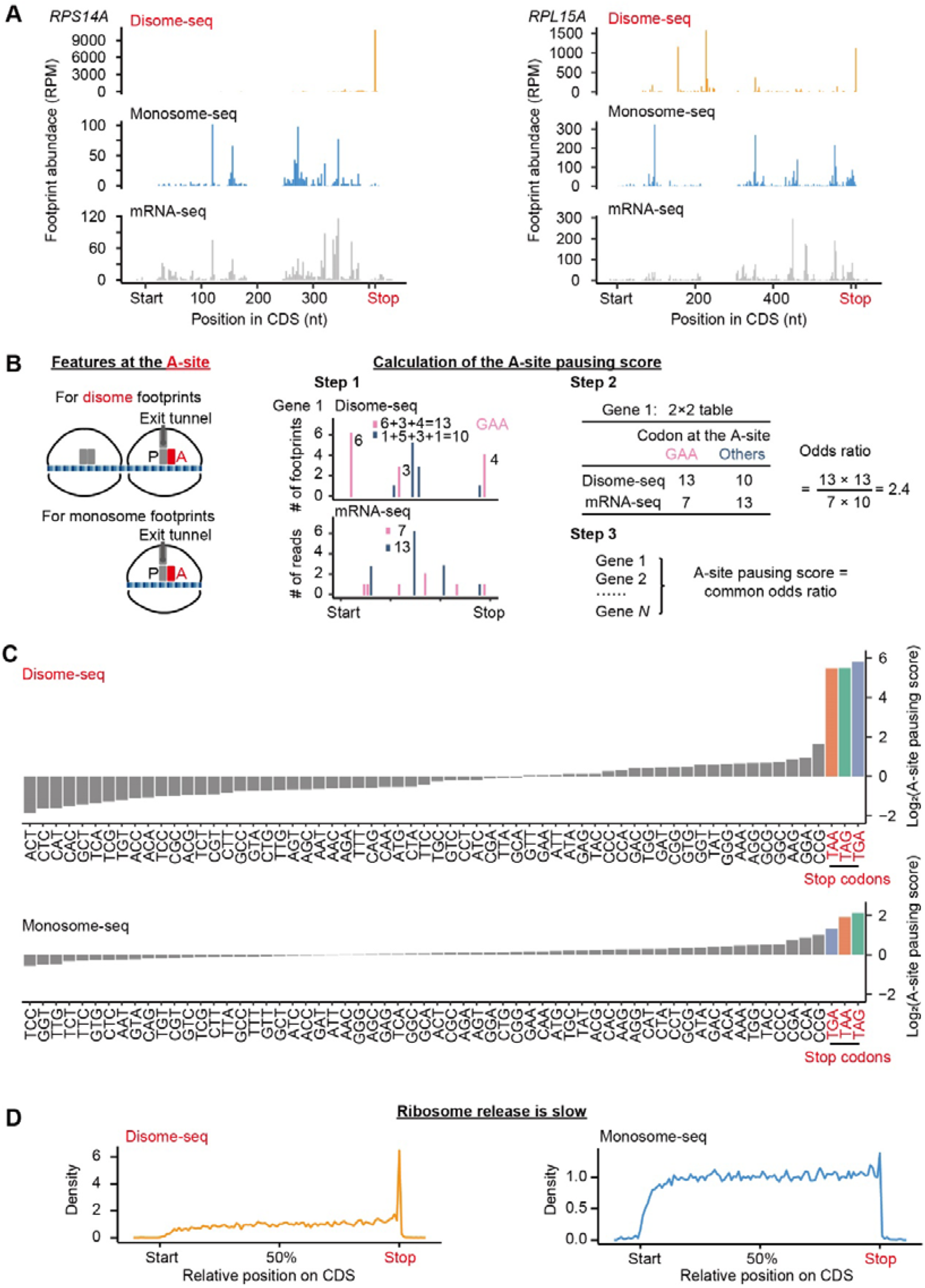
Ribosome collisions are induced by stop codons. (A) Two genes (*RPS14A* and *RPL15A*) exemplify ribosome collisions at stop codons. (B) Schematic of the calculation of the A-site pausing score. We counted the numbers of disome (or monosome) footprints with the concerning codon (GAA as an example here) or the other codons (all 63 non-GAA codons) at their A-sites, respectively. The mRNA-seq reads were used to control for the codon frequency in the transcript. The A-site pausing score was defined as the common odds ratio among genes, and the *P*-value was given by the Mantel-Haenszel test. (C) The A-site pausing scores in ascending order for disome and monosome footprints, respectively. (D) Aggregated profiles of footprint densities over 3527 (disome-seq, in orange) and 5230 (monosome-seq, in blue) genes, aligned in the CDS, are shown.

All three stop codons exhibited extremely high A-site pausing scores in disome-seq while their A-site pausing scores in monosome-seq were only a bit higher than the amino-acid coding codons’ (**Figure 3C**). The results were consistent between biological replicates (**Figure S4A**). To avoid artifacts generated by PCR amplification bias during the preparation of the high-throughput sequencing library, we used UMI to exclude PCR duplicates. Three stop codons still exhibited extremely high A-site pausing scores (**Figure S4B**). Consistently, disome reads accumulated at stop codons at the genomic scale (**Figure 3D**). Collectively, these observations suggest that ribosome release is slow.

### Ribosome collisions occur preferentially at the amino acids that terminate α-helices

To determine if peptide-bond formation causes ribosome collisions we similarly calculated the P-site pausing score for each of the 20 amino acids. Five amino acids (e.g., proline, glycine, asparagine, cysteine, and lysine) showed significantly > 1 P-site pausing scores in disome-seq (**Figure 4A**). Among them, proline and glycine are poor substrates for the formation of a peptide bond (Rodnina, 2016) and therefore, slow down translation elongation and lead to ribosome collisions. It remains unclear how other amino acids induce ribosome collisions. Nevertheless, these amino acids share the same property that they weaken the stability of the α-helix, a prominent secondary structure of proteins (Nelson et al., 2008). Proline and glycine are conformationally too inflexible and too flexible, respectively, to form an α-helix. The bulk and shape of asparagine and cysteine also destabilize α-helices. Positively charged residues (e.g., polylysine) repel each other and also prevent the formation of an α-helix (Nelson et al., 2008). The P-site pausing score and the propensity to terminate an α-helical conformation (Bryson et al., 1995; Myers et al., 1997) were positively correlated among 20 amino acids in disome-seq (*r* = 0.56, **Figure 4B**) but not in the monosome-seq (*r* = 0.21, **Figure 4B**), suggesting a sequence-mediated coupling between ribosome collisions and the folding of α-helices — a translational pause occurring right after the translational completion of an α-helix may benefit the cell by avoiding the folding interference from the downstream peptides.

**Figure 4.**
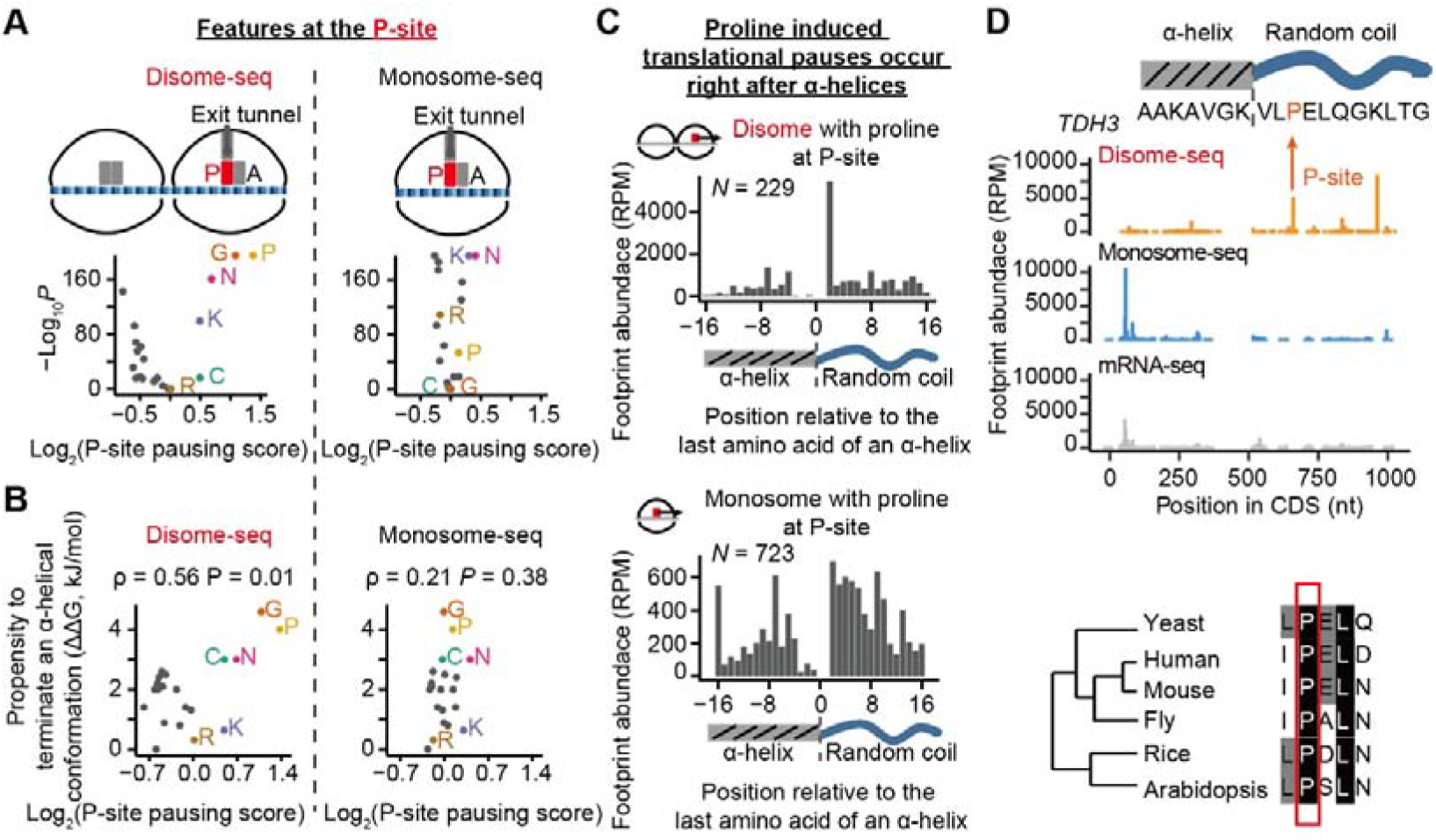
Ribosome collisions occur at the amino acids that terminateα-helixes. (A) Volcano plot shows the P-site pausing score (based on disome and monosome footprints, respectively) of each amino acid and the corresponding *P*-value in the Mantel-Haenszel test. Six amino acids with > 1 P-site pausing scores in disome-seq are labeled. P, proline; G, glycine; N, asparagine; K, lysine; C, cysteine; R, arginine. (B) The propensity to terminate an α-helix of an amino acid (ΔG relative to alanine, in the unit of kJ/mol) is plotted against its P-site pausing score. The *P*-value was given by Pearson’s correlation. (C) Aggregated profiles of disome footprint (A-site of the 3′-ribosome) and monosome footprint (A-site) frequencies are plotted against protein position and aligned by the last amino acid of an α-helix (set at position 0). (D) A ribosome collision occurs right after an α-helix of Tdh3p (top). The proline is conserved across eukaryotic kingdoms (bottom).

Such coupling can be observed at the genomic scale — disomes induced by prolines frequently took place right after α-helices (**Figure 4C**), as well as in individual genes (*TDH3* as an example in **Figure 4D**), but was not observed in monosome-seq (**Figure 4C-D**). Furthermore, such pausing signals were often evolutionarily conserved despite being within intrinsically disordered regions that are often evolutionarily dynamic (Brown et al., 2010). For example, the disome-inducing proline in Tdh3p was conserved across eukaryotic species (**Figure 4D**).

### Ribosome collisions are promoted by polylysine in the exit tunnel

Positively charged amino acids can hinder translation elongation by interacting with the negatively charged exit tunnel (Charneski and Hurst, 2013). To test this hypothesis with disome-seq data, we calculated the exit-tunnel pausing score from the enrichment of each of the (20^3^ =) 8000 amino-acid 3-mers in the 20-amino-acid region upstream of the P-site of the 3′-ribosome, with the corresponding 3-mer frequency in the transcript as a control. Ribosome collisions were strongly associated with positively charged 3-mers (e.g., RKR) when in exit tunnel; by contrast, such pausing signals were obscured in monosome-seq (**Figure 5A**). Triple-lysine had the strongest collision signal (**Figure 5A**), which was mainly contributed by triple-AAG (**Figure S5**).

**Figure 5.**
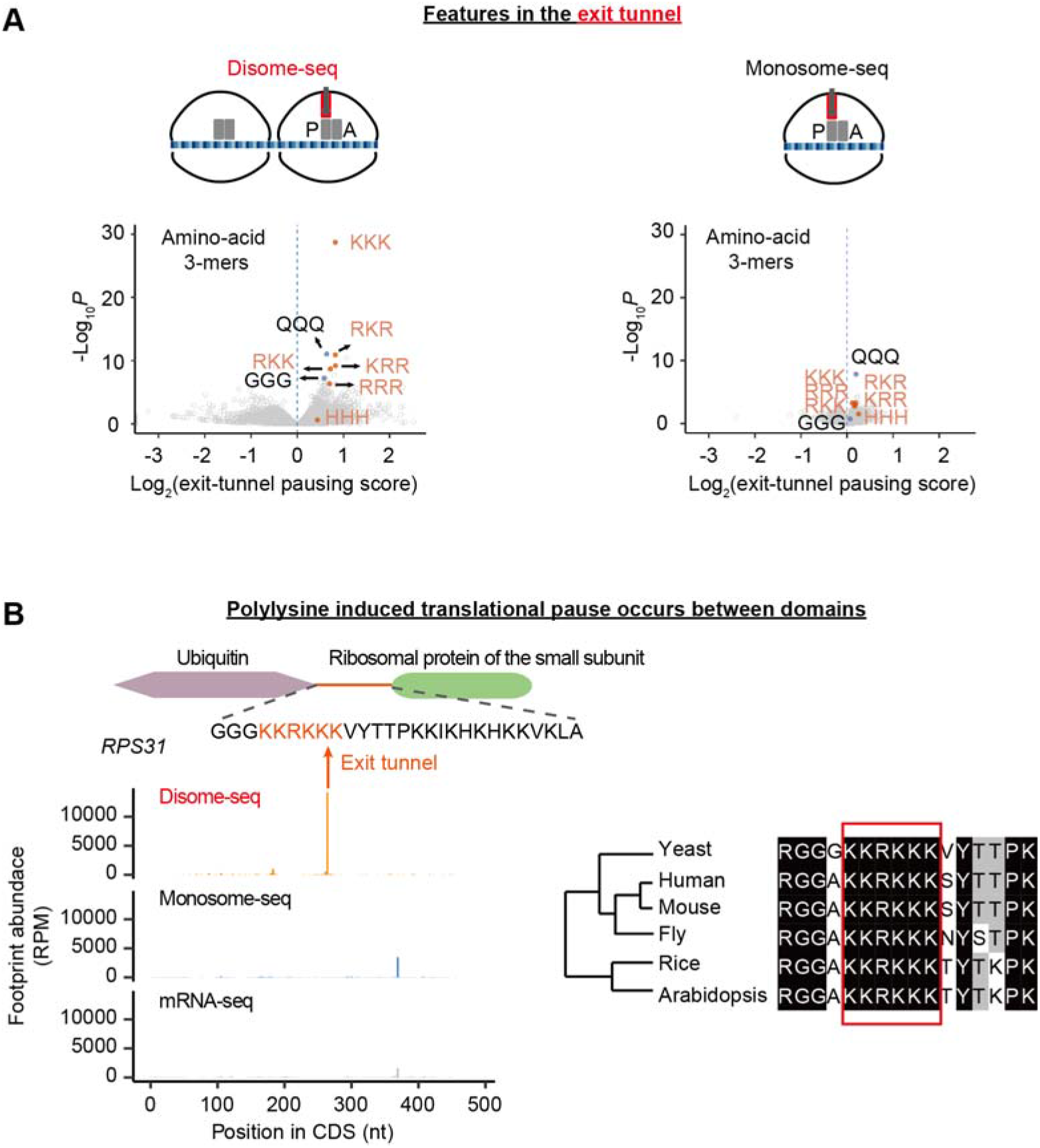
Ribosome collisions are induced by polylysine in the exit tunnel. (A) Volcano plot shows the exit-tunnel pausing score (based on disome and monosome footprints, respectively) of each of the 8000 amino-acid 3-mers and the corresponding *P*-value given by the Mantel-Haenszel test. (B) A ribosome collision induced by polylysine occurs in the linker region between two domains of RPS31p (left). The polylysine (plus an arginine) sequence is conserved across eukaryotic kingdoms (right).

The lysine tract often appears in the linker regions of domains because the repellence among the positively charged residues prevents the formation of a domain (Nelson et al., 2008). As an example, polylysine-induced ribosome collision occurs in the linker between the two domains of RPS31p, the fusion protein of ubiquitin and ribosomal protein (**Figure 5B**), raising the possibility that a translational pause benefits the cell by providing sufficient time for the co-translational folding of the upstream domain. Consistently, despite being in an intrinsically disordered region, these positively charged amino acids (five lysines and one arginine) are conserved across eukaryotes (**Figure 5B**).

### The translation of the majority of disomes is likely to resume

The pervasiveness of ribosome collisions detected in fast-proliferating cells (**Figure 2**) and the ubiquitous sequence signals of disomes (e.g., proline and polylysine, **Figures 4**–**5**) in endogenous genes is astonishing since the di-ribosome observed previously in heterologous genes could lead to the decay of both mRNA and nascent peptides through the RQC pathway (Joazeiro, 2017). To test if the disomes collected from endogenous genes in fast-proliferating cells are structurally identical to the RQC-inducing di-ribosomes (Ikeuchi et al., 2019), we performed cryo-EM analyses of disomes. It showed a structure of two ribosomes linked by a bent mRNA, and therefore, two 40S subunits oriented towards each other (**Figure 6A**). Furthermore, the 5′-colliding ribosomes were detected in the rotate state that harbors hybrid A/P and P/E tRNAs (**Figure 6A**), indicating that they cannot further translocate, likely due to the road-blocking 3′-leading ribosomes. These observations are consistent with the structure of di-ribosomes induced by a strong stalling signal in cell-free systems (Ikeuchi et al., 2019).

**Figure 6.**
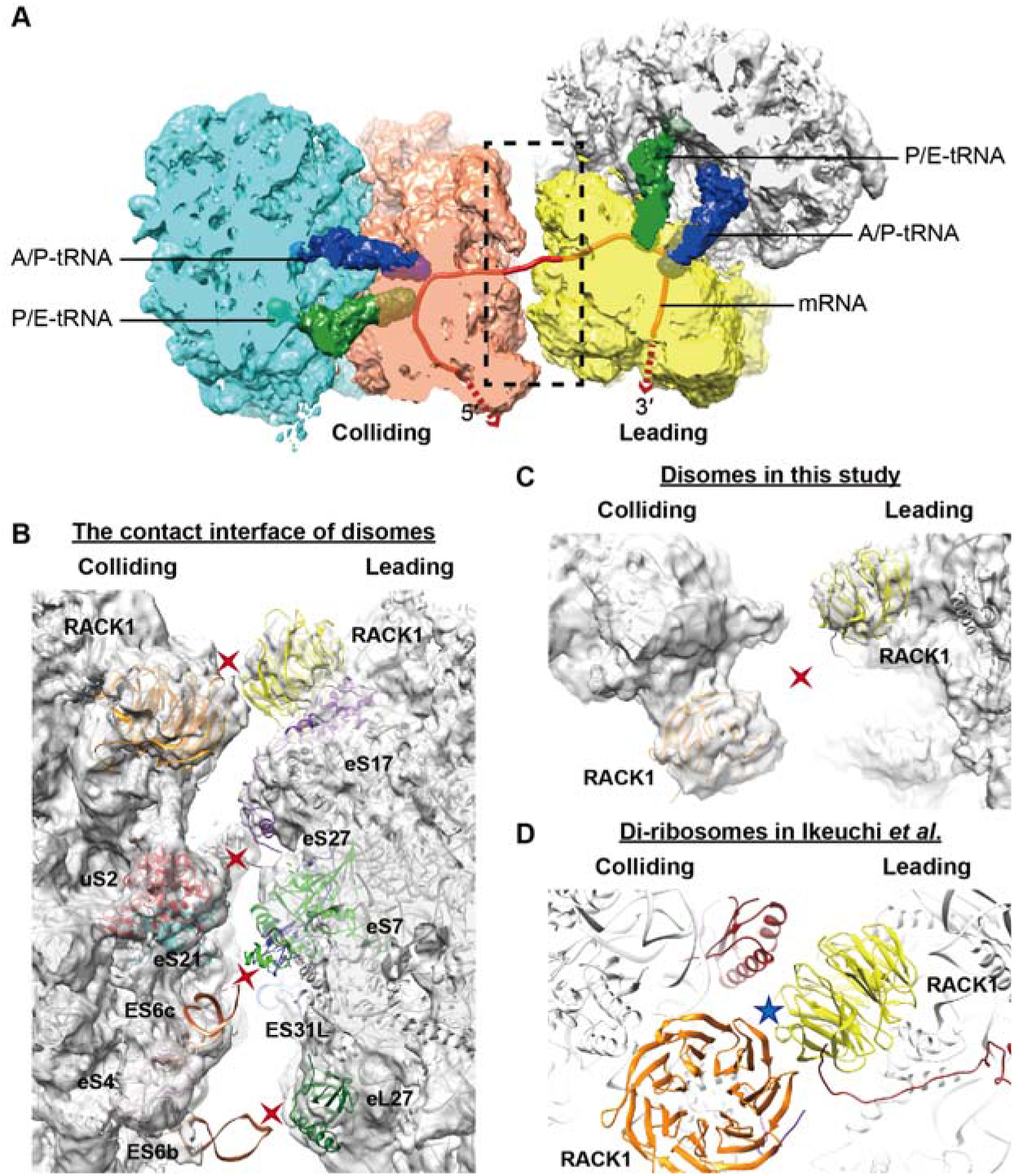
The cryo-EM structure of disomes. (A) The top view of the cryo-EM reconstruction of disomes. The 3′-ribosome is referred to as the leading ribosome, and the 5′-ribosome is referred to as the colliding ribosome. Both leading and colliding ribosomes are in the rotate state with hybrid A/P (blue) and P/E tRNAs (green). mRNA is highlighted in red. (B) The zoomed-in detail of the contact interface between the 40S subunits of the leading and colliding ribosomes of a disome. (C-D) The zoomed-in view of the 40S head-to-head contact site of a disome (C) and that of the di-ribosome structure reported in Ikeuchi *et al.*. (D) The blue star indicates a strong interaction between the two ribosomes in di-ribosomes, while red crosses indicate the interactions absent in disomes.

However, two distinct features indicate the essential difference between the disomes and di-ribosomes. First, the connection between two ribosomes was relatively flexible in disomes (**Figure 6B–C** and **Figures S6**), in contrast to the rigid one detected in di-ribosomes. In di-ribosomes, a large and tight interface between its composing ribosomes was formed, which was considered recognizable by the RQC initiation factors such as Hel2p (Ikeuchi et al., 2019; Matsuo et al., 2017; Sundaramoorthy et al., 2017). However, the ribosome interface is much weaker or nearly absent in our structure of the disome due to a twist between two ribosomes, suggesting that they may not be recognizable by Hel2p to trigger RQC (**Figure 6B–C** and **Figures S6**). For instance, a tight interaction was seen between the two RACK1 (Asc1p) — the protein involved in the initiation of RQC — from the two ribosomes in the di-ribosome (**Figure 6D**), but it was completely lost in our structure (**Figure 6C**). Second, the 3′-leading ribosome of disomes was detected in the rotate state (**Figure 6A**), in sharp contrast to the POST state (with P- and E-site tRNAs) of the 3′-leading ribosome in di-ribosomes, a particular stalling state adopted by the stalled 80S ribosomes before collision (Ikeuchi et al., 2019). The rotate state of the 3′-leading ribosome in disomes indicates their transient-pausing state and the capability to continue translation. Collectively, the results of cryo-EM analyses are consistent with the idea that disomes represent snapshots of translocation-competent ribosomes that do not trigger RQC.

### Ribosome collisions facilitate the recruitment of chaperones

To understand the function of ribosome collisions, we attempted to identify disome-specific ribosome components. We labeled ribosome proteins with stable isotopes, digested the mRNA with RNase I, separated disomes from monosomes with sucrose density gradient centrifugation, mixed heavy-labeled disomes with light-labeled monosomes (or light-labeled disomes with heavy-labeled monosomes in a label-swap replicate), and performed tandem mass spectrometry (MS/MS, **Figure 7A**). In addition to ribosomal proteins, some highly expressed metabolic enzymes were also identified (**Figure 7B**, **Figure S7A**, and **Tables S4**, **S5**); we speculated that they were nascent peptides of incomplete protein translation attached to the ribosomes. Consistently, the peptides of these enzymes captured by MS/MS tended to be in the first half of coding sequences (common odds ratio = 1.9, *P* = 0.001, the Mantel-Haenszel test). For example, all detected peptides of FAS2p, a fatty acid synthetase, were in the first half (**Figure S7B**). By contrast, the peptides of ribosomal proteins captured by MS/MS were evenly distributed in the first and second half of coding sequences (common odds ratio = 1.0, *P* = 0.7).

**Figure 7.**
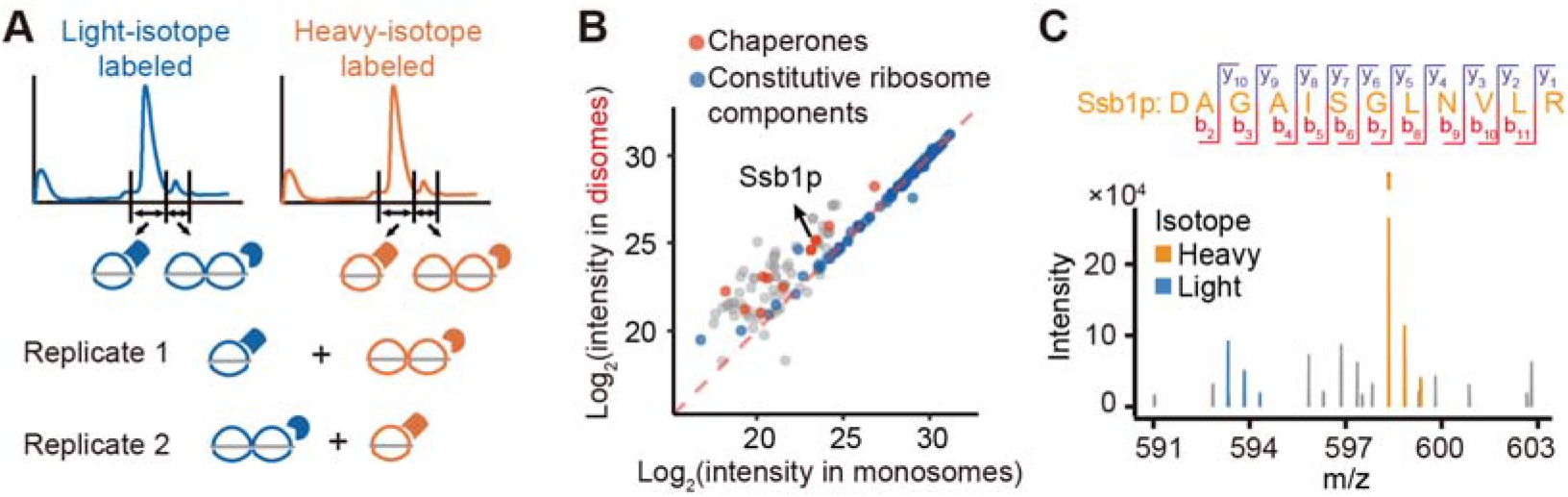
Ribosome collisions recruit chaperones. (A) The schematic of the detection of disome enriched proteins through SILAC. (B) The protein intensities in disomes were plotted against those in monosomes. Chaperones (orange) are enriched in disomes. Most constitutive ribosomal proteins (blue) on diagonal are evenly distributed between monosomes and disomes. (C) The m/z (mass divided by charge number) spectrum of Ssb1p and its fragments detected in MS/MS.

Most proteins have similar abundance in disomes and monosomes and therefore, were on the diagonal in the scatter plot showing protein intensities in disomes vs. monosomes; many were the constitutive ribosome components (**Figure 7B** and **Figure S7A**). Proteins above the diagonal line were the disome-enriched proteins (**Figure 7B**, **Figure S7A**, **Tables S4**, and **Tables S5**). Proteins recruited by RQC such as Hel2p, Slh1p, Cue3p, and Rqt4p (Ikeuchi et al., 2018; Ikeuchi et al., 2019; Juszkiewicz et al., 2018) were not observed significantly enriched in the disome fraction (**Tables S4** and **S5**). However, all 11 chaperones identified by MS/MS had > 1.5-fold abundance in disomes than in monosomes (*P* = 1×10^−6^, Fisher’s exact test, **Figure 7B**; *P* = 1×10^−4^, **Figure S7A**). For example, although SSB1p was reported as a ribosome-associated chaperone (Doring et al., 2017), how SSB1p is distributed among ribosomes remained unclear. We showed that SSB1p was associated primarily with disomes (**Figure 7B-C**), suggesting that ribosome collisions promote the recruitment of chaperones to aid in the folding of nascent peptides.

## DISCUSSION

Translation elongation plays a central role in regulating protein folding. The key to studying such regulation is to unveil the dynamics of ribosomes during translation. Traditional ribo-seq (i.e., monosome-seq) solves the problem to some extent but misses collided ribosomes, which are present in 75% of genes in yeast and provide unique information about translational pausing. In this study, we characterize the genomic landscape of disome footprints, which enables us to identify the leading sequence features regulating translation elongation, and provide a mechanism that chaperones sense translation elongation rate through ribosome collisions to determine which proteins require co-translational folding.

It is worth noting that disome footprints are fundamentally different from the transcripts binding to two ribosomes, in which two ribosomes are usually sparsely distributed, and the mRNA fragment between them is sensitive to RNase I. To illustrate this point experimentally, we collected the transcripts binding to two ribosomes, performed RNase I digestion, and sequenced ribosome protected mRNA fragments (**Figure S8A**). Not unexpectedly, the footprints repeatedly observed in disome-seq did not show up in such experiments (**Figure S8B**).

In addition to the 58-nt disome footprints whose conformation was resolved in **Figure 1D**, the 5′-end of the 61-nt fragments also exhibited an apparent 3-nt periodicity (**Figure S3B, D**), suggesting the presence of a second disome conformation. The disomes accumulated at the stop codons (**Figure S9A**) indicate that the 61-nt disome footprints were composed of two 28-nt monosome footprints spaced by five nucleotides. Although in principle the 5′-ribosome in such disomes can make another move, two ribosomes are sufficiently close to each other in such quasi-collision that the mRNA fragment between them is at least partially resistant to RNase I digestion. The P-site pausing scores estimated from 58 or 61-nt disome footprints were highly correlated (ρ = 0.88, *P* = 1.2×10^−6^, Spearman’s correlation, **Figure S9B**), indicating that the 61-nt quasi-collisions also reflect translational pauses. Therefore, they were combined with the 58-nt footprints for the analyses of this study.

In addition to disomes, translational pauses may further lead to the collision of three stacked ribosomes — trisomes (Pelechano et al., 2015). Indeed, after RNase I digestion, a small number of particles were observed at the ~ 62 mm position after the sucrose gradient centrifugation (**Figure 1B**); the density of these particles was higher than disomes and was similar to transcripts binding to three ribosomes, suggesting that they were trisomes. Although the abundance of these trisomes was insufficient for a trisome-seq experiment, the presence of trisomes upstream of stop codons was evident in disome-seq by the footprints with the 5′-end ~75-nt upstream of the stop codon (**Figure S9C**).

The disomes detected in fast proliferating cells may simply reflect the inevitable nature of stochastic ribosome collisions since an mRNA is often translated simultaneously by multiple ribosomes. To test the idea, we estimated the collision propensity of each gene by calculating the number of ribosome collisions per mRNA per unit length and correlated this propensity with ribosome density of the gene (approximated by the number of monosomes per mRNA per unit length). Consistent with this idea, collision propensity was positively correlated with ribosome density (ρ = 0.35, *P* < 10^−100^, **Figure S10A**) — the 3′-pausing ribosomes are easier to be caught up by a 5′-ribosomes when mRNA is crowded with more ribosomes. In these disomes, translation is transiently interrupted and to be continued. Consistently, ribosome density was lower downstream of disomes (**Figure S10B**), which is easy to understand with a metaphor — fewer vehicles are on the highway after passing a traffic jam.

It remains possible that a population of disomes may trigger the RQC pathway; however, such disomes may have been quickly disassembled and therefore, are hardly detected by disome-seq in the wild-type yeast. Nevertheless, such disomes will accumulate in the mutants of the RQC pathway (*hel2* for example). Our disome-seq results provide a genomic landscape of disomes in the wild-type, which will facilitate the identification of endogenous signals of the RQC pathways in the future.

In addition to the sequence features identified at the A-site, P-site, and exit tunnel, others may also play a role. For example, mRNA secondary structure has been shown to cause translational pausing (Yang et al., 2014), which was also detected in this study; mRNA downstream of disomes exhibited stronger secondary structure (**Figure S10C**). Relatedly, RNA helicases TIF1p and TIF2p were recruited by disomes (**Tables S4**, **S5**), potentially removing mRNA secondary structures to relieve translational pauses. There are also sequence features that lead to ribosome collisions with unknown mechanisms. For example, two collision-inducing 3-mers, QQQ and GGG, were identified in the exit tunnel (**Figure 5A**); they are not positively charged. Nevertheless, both are related to protein folding to some extent. The former is a well-known signal for protein misfolding (Williams and Paulson, 2008), and the latter is a prominent signal for intrinsic protein disorder (Rauscher et al., 2006).

Yeast genes often use non-optimal codons at the 5′ of the coding sequences, which has been speculated to serve as a “ramp” (Tuller et al., 2010). However, we did not detect any accumulation of ribosomes downstream of the start codon in either monosome-seq or disome-seq in this study (**Figure 3D**). It is likely that the previously observed accumulation of ribosomes around the start codon (Ingolia et al., 2009) is a byproduct of the pre-treatment with cycloheximide (Hussmann et al., 2015; Pelechano et al., 2015; Weinberg et al., 2016) that partially inhibits translation elongation but does not block initiation (Schneider-Poetsch et al., 2010).

It is apparently costly when ribosomes are sequestered upstream of the stop codon waiting for release since they are not used for active protein synthesis. Nevertheless, slow ribosome release may benefit the cell by providing the newly synthesized peptide sufficient time to fold within the exit tunnel of ribosomes (Nilsson et al., 2015), rather than in the complex cytoplasmic environment. Some amino-acid sequences (e.g., proline and glycine at the P-site as well as polylysine in the exit tunnel) induce translational pauses as well (**Figures 4** and **5**); they are also signals for the termination of α-helices during protein folding (**Figure 4B**). These amino-acid sequences may provide the time and subcellular environments for the folding of the newly synthesized peptide inside the exit tunnel, especially α-helices. The constriction site of exit tunnel, the narrowest region of the exit tunnel, is ~ 10 Å in width (Fedyukina and Cavagnero, 2011), sufficient for co-translational folded α-helices (~ 5 Å in width) (Voss et al., 2006) to pass by.

Such dual role of amino acids can catalyze the evolution of protein structures because it does not take additional time for placing a translational pausing signal after an evolutionary change in protein structure — the amino-acid substitution that results in an innovation in protein structure on the same time confers a translational pause, making the emergent protein structure ready to be folded. Together with the chaperones recruited co-translationally by collided ribosomes, the correct folding of a novel protein structure is warranted during evolution.

## METHODS

### Polysome profiling and disome-seq

The laboratory strain BY4742 (*MATα his3Δ1 leu2Δ0 lys2Δ0 uraΔ0*) was cultivated at 30°C in the rich medium YPD (1% yeast extract, 2% peptone, and 2% dextrose) or in the SC−His+3-AT medium (synthetic complete medium with histidine dropped-out and 100 mM 3-AT added). We harvested cells following previous studies (Guydosh and Green, 2014). Briefly, in the absence of cycloheximide co-culture, cells were collected at OD_660_ ~0.6 by vacuum filtration and were immediately frozen in liquid nitrogen. Ribosomes were extracted with the polysome lysis buffer (PLB), which contained 200 mM Tris-HCl (pH 8.0), 200 mM KCl, 35 mM MgCl_2_, 1% (v/v) Triton X-100, 5 mM DTT, and 50 μg/mL cycloheximide.

For the polysome profiling experiments, the extracted ribosomes were pelleted through a 30 mL sucrose cushion containing 400 mM Tris-HCl (pH 8.0), 200 mM KCl, 30 mM MgCl_2_, 1.75 M sucrose, 5 mM DTT, and 50 μg/mL cycloheximide, by ultracentrifugation at 4°C overnight (33500 rpm, Beckman, 70Ti rotor). The ribosome pellet was dissolved in 300 μL resuspension buffer (20 mM Tris-HCl pH 8.0, 140 mM KCl, 5 mM MgCl_2_, and 50 μg/mL cycloheximide), and separated by ultracentrifugation at 4°C for 3 hours (35300 rpm, Beckman, SW41 rotor) through a 5-50% sucrose gradient (40 mM Tris-HCl pH 8.4, 20 mM KCl, 10 mM MgCl_2_, and 50 μg/mL cycloheximide) prepared by Gradient Master (Biocomp). The profiling signals were recorded by Piston Gradient Fractionator (Biocomp).

For the disome-seq experiments, 50000 A260 units of ribosome dissolved in the PLB buffer were treated with 750 U RNase I (Ambion, AM2294) at 25°C for 2 hours. RNA was extracted from the enzyme reaction system with hot phenol and was separated on a 17% (w/v) 7 M urea denaturing polyacrylamide gel in a 0.5×Tris-borate-EDTA (TBE) electrophoresis buffer. RNA fragments with the length of 25-30 nts or 50-80 nts were extracted by gel crushing and further incubated with an RNA gel extraction buffer (300 mM NaOAc pH 5.2, 10 mM Tris-HCl pH 8.0, 1 mM EDTA pH 8.0) overnight. The purified RNA fragments were subjected to small RNA library construction for Illumina sequencing (Gnomegen, k02420). The 5′-RNA adaptor contained a 3-nt random sequence at the 3′-end to avoid biased ligation. Monosome and disome footprints were sequenced with single-end 50 and paired-end 100 modes on BGISEQ-500 (BGI Group), respectively.

### Mapping reads to the yeast genome

The 3-nt random sequence at the 5′-end of each sequencing read was removed. The removed 3-nt sequence was added to the head of each read in the fastq format as the UMI, which can serve to remove PCR duplications generated during Illumina library preparation. The sequence identical to the 3′-sequencing adaptor was also trimmed in each read using cutadapt V1.16 (http://gensoft.pasteur.fr/docs/cutadapt/1.6/index.html). Reads without 3′-adaptor sequence were removed since they (> 46 nts for monosome-seq or > 96 nts for disome-seq) were much longer than the expectation (25-30 nts for monosome-seq and 50-80 nts for disome-seq). Trimmed reads that were shorter than 20 nts were also excluded for further analyses.

The *S. cerevisiae* genome (SGD R64-1-1) (https://www.yeastgenome.org/) was used as the reference. The trimmed reads were mapped against rRNA with bowtie V1.2.2 (Langmead et al., 2009) (http://bowtie-bio.sourceforge.net/manual.shtml); the mapped reads were filtered to avoid rRNA contamination. The rest reads were aligned against coding sequences with the--no-novel-juncs parameter using Tophat V2.1.1 (Kim et al., 2013) (https://ccb.jhu.edu/software/tophat/manual.shtml). Reads with multiple alignments or with mapping quality < 30 were discarded. Biological replicates were highly correlated (**Figure S2**) and were combined in the majority of our analyses. To remove PCR duplicates generated during Illumina library preparation, sequencing reads of the same length, sequence, and UMI were counted only once. The disomes reads obtained in this study (**Table S2**) are comparable to those in previous studies (Guydosh and Green, 2014).

### The definition of the A-site

For the 58-nt disome footprints, if the 5′-end of a read was mapped to the first nucleotide of a codon (in frame), the 46^th^-48^th^ nts were defined as the A-site of the 3′-ribosome; if mapped to +1 (or +2) frame, the 45^th^-47^th^ (or 47^th^-49^th^) nts were defined as the A-site since the footprint likely shifted by +1 and −1 nt during RNase I digestion. The A-site of the 3′-ribosome was also defined for the 59-nt disome footprints: if the 5′-end was in frame, meaning that one more nt was kept at the 3′-end during RNase I digestion, the 46^th^-48^th^ nts was the A-site. If mapped to +2 (or +1) frame, meaning that one more nt was kept at the 5′-end (or 3′-end) during RNase I digestion, the 47^th^-49^th^ (or 45^th^-47^th^) nts were the A-site. For 61-nt and 62-nt disome footprints, the A-site was defined assuming that an extra codon was included in the space between two ribosomes. For the 28 and 29-nt monosome footprints, the definition of the A-site was the same as the 58 and 59-nt disome footprints except for a 30-nt offset.

### The calculation of the pausing scores

The A-site pausing score of each of the 64 codons was defined as the common odds ratio among genes calculated by the Mantel-Haenszel test. A 2×2 contingency table was generated for each gene. Disome footprints were divided into two categories, the concerning codon at the A-site of the 3′-ribosome and others. mRNA reads were used as the background to control for codon frequency in the gene; the 16^th^-18^th^ nts of a 28-nt mRNA-seq read was considered as the A-site. The P-site pausing score was estimated similarly, with the amino acid at the P-site under consideration. The exit-tunnel pausing score was estimated for each of the amino-acid 3-mers in the 20-amino-acid region upstream of the P-site; the length was conservative since peptides with 33 to 67 amino acids can fold within the exit tunnel (Lu and Deutsch, 2005). Flanking 3-mers may hitchhike on the causal 3-mer, and therefore, we combined multiple disome reads mapped to the same genomic position into one to reduce such hitchhiking effect.

Statistical analyses were performed with *R* (https://www.r-project.org/), and plots were generated with an *R* package, ggplot2 (https://ggplot2.tidyverse.org/). All statistical tests were two-sided.

### Cryo-EM

Disome purification: Ribosomes were extracted as described in the polysome profiling experiments. The disome particles were collected after a 5-50% sucrose gradient separation.

Data acquisition: Vitrified specimens were prepared by adding 4 μl disome samples at the concentration of ~ 150 nM to a glow-discharged holey carbon grid (Quantifoil R1.2/1.3), which was covered with a freshly made thin carbon film. Grids were blotted for 1 second and plunge-frozen into liquid ethane using an FEI Vitrobot Mark IV (4°C and 100% humidity). The cryo-grids were initially screened at a nominal magnification of 92000× in an FEI Tecnai Arctica microscope, equipped with an Autoloader and an acceleration voltage of 200 kV. High-quality grids were transferred to an FEI Titan Krios electron microscope operating at 300 kV, and images were collected using a K2 Summit direct electron detector (Gatan) in counting mode at a nominal magnification of ×105000, which corresponds to a pixel size of 1.373 Å at the object scale and with the defocus varying from −1.0 to −2.0 μm. The coma-free alignment was manually optimized and parallel illumination was verified before data collection. All micrographs obtained with the K2 camera were collected semi-automatically by SerialEM (Mastronarde, 2005), under low-dose conditions. Each micrograph was dose-fractionated to 32 frames with a dose rate of ~ 10.0 counts per physical pixel per second for a total exposure time of 6.4 seconds.

Data processing: Original image stacks were summed and corrected for drift and beam-induced motion at the micrograph level using MotionCor2 (Zheng et al., 2017). SPIDER (Shaikh et al., 2008) was used for micrograph screening. The contrast transfer function parameters of each micrograph were estimated by Gctf (Zhang, 2016). All 2D and 3D classification and refinement were performed with RELION-3.0 (Zivanov et al., 2018).

The disome data were initially processed using 80S monosomes as a template for particle picking. A total of 1439 micrographs were collected and 269962 particles were picked for a cascade 2D and 3D classification with a binning factor of two. Around 78% of particles were removed during several rounds of 2D and 3D classification. The remaining 59816 particles were split into five classes during the final round of 3D classification with the box size of 400 pixels (**Figure S11**). A total of 17945 particles from the two classes containing the second 80S densities were re-centered and re-extracted with a box size of 440 pixels, and were split into five classes during the final disome 3D classification. After the final round of 3D classification, a total of 4845 particles were applied for 3D auto-refine, generating a map at an overall resolution of 8.9 Å. To further improve the map to identify the conformational status of ribosomes, masked-based refinement was performed using a soft mask on leading or colliding ribosomes. In both cases the maps could be improved to 4.2 Å (gold-standard FSC 0.143 criteria), allowing the assignment of tRNA positions.

### Mass spectrometry

Stable isotope labeling with amino acids in cell culture (SILAC) was performed as described previously (Chen et al., 2019). Briefly, a strain modified from BY4742 (*MATα his3Δ1 leu2Δ0 lys2Δ0 ura3Δ0 arg4Δ0::kanMX4 car1Δ0::LEU2*) was cultured at 30°C in the regular SC medium or in the SC medium substituted with heavy isotopes (37.25 mg/L Lys8 and 20.94 mg/L Arg10). Lys8 and Arg10 represent 15N_2_^13^C_6_-lysine and ^15^N_4_^13^C_6_-arginine, respectively.

Cells were harvested at the mid-log phase (OD_660_ ~0.6), and polysomes were extracted. Polysome profiling was performed after RNase I digestion, and proteins were precipitated from the monosome or disome fraction with a double volume of 95% ethanol, respectively. The protein precipitants were dissolved with the urea buffer (8 M urea, 1 mM sodium orthovanadate, 1 mM sodium fluoride, 2.5 mM sodium pyrophosphate, 1 mM B-glycerophosphate, 0.2% tablet of protease inhibitors, and 1 mM PMSF). 150 μg light-labeled monosome protein was mixed with 150 μg heavy-labeled disome protein; 150 μg heavy-labeled monosome protein was mixed with 150 μg light-labeled disome protein as a replicate. The protein mixtures were subject to the MS/MS analysis on Orbitrap Elite (Thermo Fisher Scientific).

The raw data were processed with MaxQuant V1.5.8.3 (Cox and Mann, 2008) (https://www.biochem.mpg.de/5111795/maxquant) using default parameters. All raw data were searched against the yeast proteome with the addition of potential contaminants. The protease was set as trypsin/P and trypsin. Two missed cuts were allowed. The protein intensities were retrieved from the output file (proteinGroups.txt) of MaxQuant. The proteins recruited by ribosome collision can be identified by comparing relative abundance between disome and monosome fractions. If a significant fraction of monosomes were vacant, proteins associated with translating ribosomes could in principle also be identified associated with disomes. However, this is unlikely the case since all monosomes (100%) detected in the cryo-EM analyses accommodated tRNAs, indicating they are mostly translating ribosomes.

If the peptides of metabolic enzymes were detected in the ribosome particles because they were nascent peptides of incomplete protein translation, they should enrich in the first half (N-terminus) of a protein. To determine if it is true, we divided peptides detected into two categories, according to whether the first amino acid of the peptide belonged to the first half of the protein or not. With the theoretical digested peptides (arginine or lysine at the C terminus) as a background, a 2×2 contingency table was generated for each gene. The common odds ratio among genes was calculated by the Mantel-Haenszel test.

### The definition of α-helices

The protein secondary structure annotation of yeast protein was retrieved from UniProt (UniProt Consortium, 2019) (https://www.uniprot.org/). Adjacent α-helices without a gap were concatenated. Short α-helices composed of 4 amino acids or less were filtered. The secondary structure of Tdh3p was predicted with the online secondary structure prediction method, GOR IV (Garnier et al., 1996) at https://npsa-prabi.ibcp.fr/.

### Alignment of proteins from different species

Homologous genes were identified by BLAST (https://blast.ncbi.nlm.nih.gov/Blast.cgi). Amino-acid sequences from multiple species were aligned by Clustal X (Larkin et al., 2007). Species trees were constructed with TimeTree (Kumar et al., 2017) at http://www.timetree.org.

### Prediction of the minimum free energy of an mRNA fragment

The minimum free energy of the 30-nt mRNA downstream of a disome footprint was calculated with the RNAfold V2.4.13 of the Vienna RNA package (Lorenz et al., 2011) (http://rna.tbi.univie.ac.at/).

## ACKNOWLEDGMENTS

We thank Lucas Carey (Peking University) for commenting on and editing the manuscript. We thank Xiahe Huang and Yingchun Wang (IGDB, CAS) for technical support in proteomics, Ting Li (IGDB, CAS) in ribosome profiling, Rongxin Yang (IGDB, CAS) in molecular biological experiments, Shaohuan Wu (IGDB, CAS) and Haotian Guo (Huazhong University of Science and Technology) in computational biology. We thank Jinzhong Lin (Fudan University) and Xiaofeng Cao (IGDB, CAS) for discussion. We also thank Addgene (http://www.addgene.org/) for providing plasmids. This work was supported by grants from the National Key R&D Program of China (2019YFA0508700) and the National Natural Science Foundation of China (31900455 to T.Z., 31725007 and 31630087 to N.G., and 91331112 to W.Q.).

## AUTHOR CONTRIBUTIONS

T.Z. and W.Q. designed the experiments. T.Z. performed disome-seq, mRNA-seq, and molecular biological experiments. T.Z. and J.W. performed ribo-seq. T.Z. and S.C. prepared the samples for MS/MS. Y.-M.C. and T.Z. performed data analyses for the above-mentioned experiments. Y.L., T.Z., and N.G. performed cryo-EM and the corresponding data analyses. T.Z., Y.-M.C., and W.Q. wrote the manuscript.

## DATA AVAILABILITY

The accession number for the high-throughput sequencing data reported in this paper is GSA: PRJCA001692 (http://bigd.big.ac.cn/gsa) (Wang et al., 2017). The accession number for the mass spectrometry proteomics data reported in this paper is ProteomeXchange: PXD015271 (http://proteomecentral.proteomexchange.org) (Perez-Riverol et al., 2019). Figures 1-5 are associated with raw sequencing data, Figure 6 is associated with cryo-EM data, and Figure 7 is associated with raw proteomics data.

## CODE AVAILABILITY

Codes to analyze the data are available at GitHub (https://github.com/mingming-cgz/Disome-seq).

