## Supplementary materials for "Disome-seq reveals widespread ribosome collisions that recruit co-translational chaperones"

Supplementary information includes Supplementary Figures S1-11 and Supplementary Tables S1-5.

**Supplementary Figures**

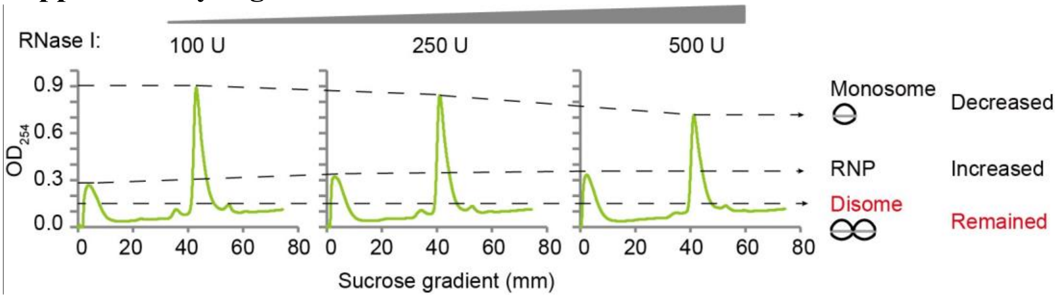


**Figure S1. Disomes persisted after RNase digestion.**

Samples containing an equal amount of ribosome-bound mRNA (5000 A260 unit) were treated with 100 U, 250 U, and 500 U RNase I, respectively. As the concentration of RNase I increased, the abundance of monosome reduced, and that of free ribonucleoprotein (RNP) increased, suggesting the disruption of ribosomes by the excessive RNase I digestion. However, the disome persisted – the mRNA fragment in-between was resistant to the RNase digestion likely due to the steric effect.

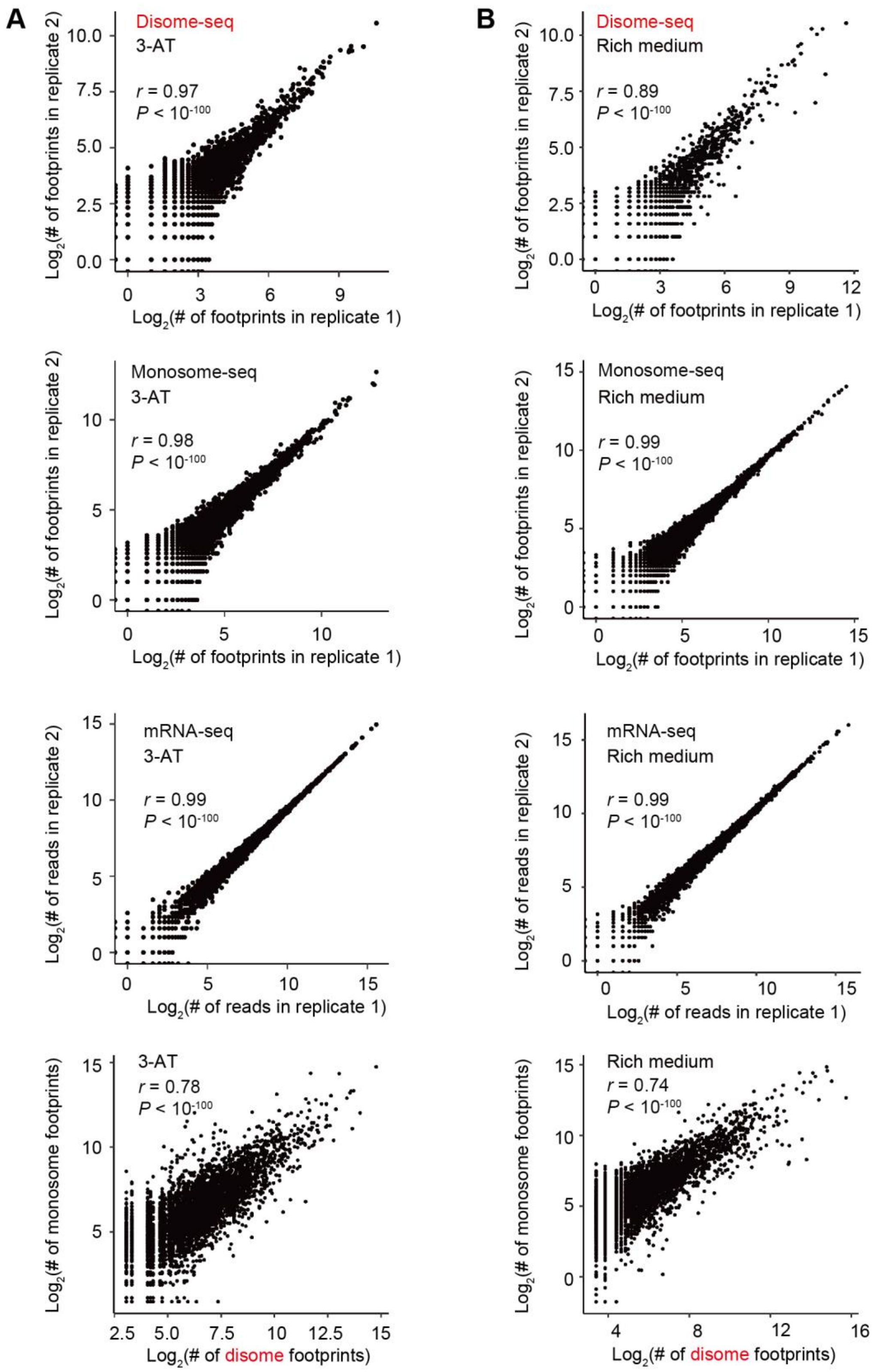


**Figure S2. Correlations between libraries of disome-seq, monosome-seq, and mRNA-seq.**

(A) The top three charts are the scatter plots of the number of mapped reads in each gene between biological replicates. The bottom chart shows the correlation of reads in each gene between disome-seq and monosome-seq, in the unit of reads per million.

The average of two replicates is shown. All libraries were obtained from 3-AT treated yeast cells. Each dot represents a gene. The *P*-values were given by Pearson’s correlation.

(B) Same to (A), except yeast cells were cultured in the rich medium.

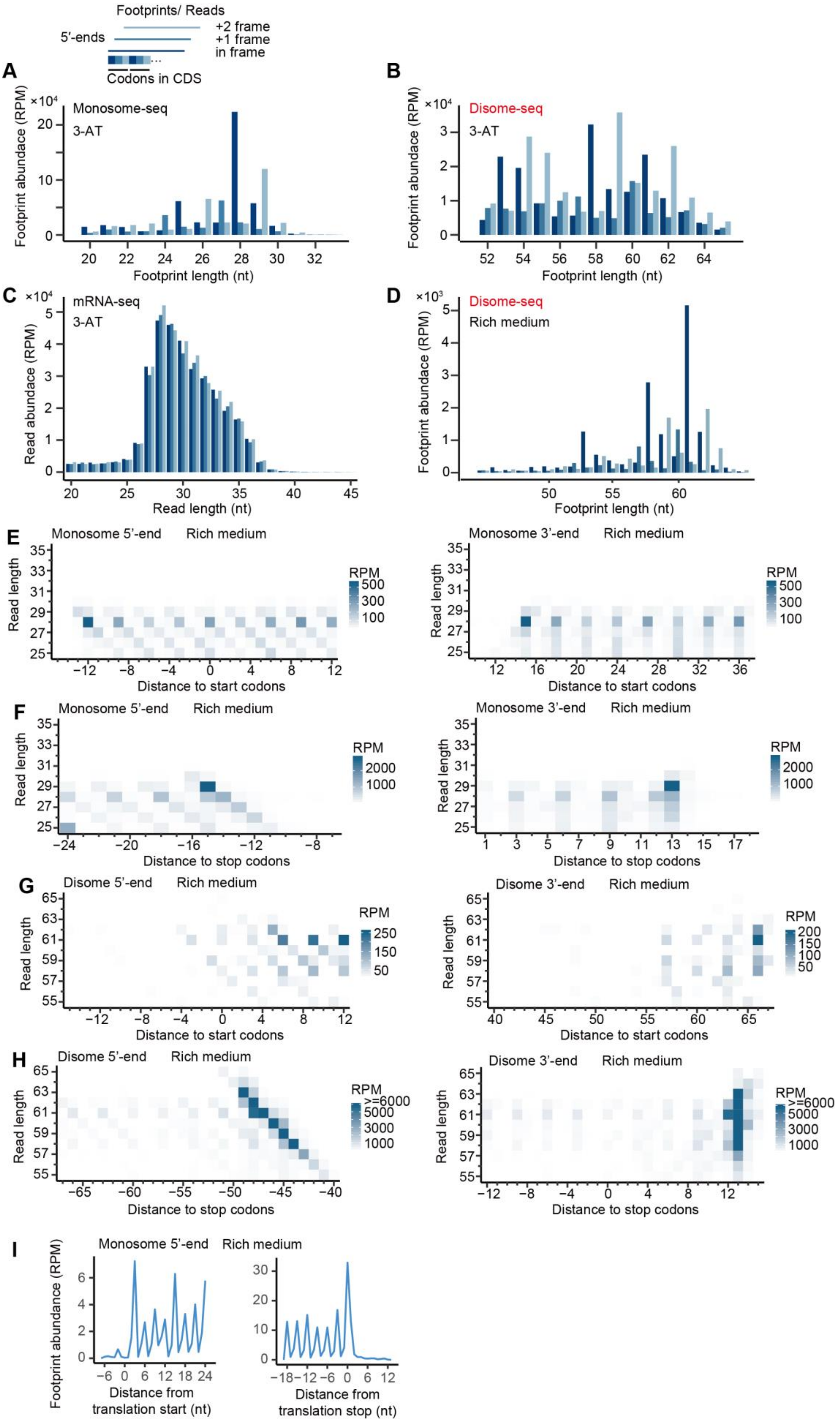

Footprints/ Reads
5′-ends
+2 frame
+1 frame
in frame
Codons in CDS
A
Monosome-seq
3-AT
Footprint abundance (RPM)
Footprint length (nt)
B
Disome-seq
3-AT
Footprint abundance (RPM)
Footprint length (nt)
C
mRNA-seq
3-AT
Read abundace (RPM)
Read length (nt)
D
Disome-seq
Rich medium
Footprint abundance (RPM)
Footprint length (nt)
E
Monosome 5'-end Rich medium
Monosome 3'-end Rich medium
Read length
RPM
Distance to start codons
F
Monosome 5'-end Rich medium
Monosome 3'-end Rich medium
Read length
RPM
Distance to stop codons
G
Disome 5'-end Rich medium
Disome 3'-end Rich medium
Read length
RPM
Distance to start codons
H
Disome 5'-end Rich medium
Disome 3'-end Rich medium
Read length
RPM
>=6000
Distance to stop codons
I
Monosome 5'-end Rich medium
Footprint abundance (RPM)
Distance from translation start (nt)
Distance from translation stop (nt)

**Figure S3. The size distribution of monosome and disome footprints.**

(A-B) We mapped the monosome (A) and disome (B) footprints obtained from 3-AT treated yeast cells to the yeast genome. The 5′-end of the 28-nt monosome footprints displayed an apparent 3-nt periodicity, similar to previous observations by Ingolia *et al.* (2009). It is consistent with the non-overlapping 3-nt genetic code – ribosome moves three nucleotides per step during translation. Notably, the 5′-end of the 58-nt disome footprints displayed a similar 3-nt periodicity (B), indicating that a 58-nt disome footprint likely contained a 28-nt monosome footprint at the 5′-end. The average of two replicates is shown.

(C) The size distribution of mRNA-seq reads obtained from 3-AT treated yeast cells is shown. The mRNA-seq reads display no apparent periodicity. The average of two replicates is shown.

(D) The disome footprints obtained from yeast cells cultivated in the rich medium display an apparent 3-nt periodicity. The average of two replicates is shown.

(E-F) The monosome footprints obtained from rich medium cultivated yeast cells. The aggregated abundance profile over the start codon (E) and the stop codon (F) region. The 5′-end (left) or the 3′-end (right) position of the reads are plotted against the read length.

(G-H) Same to (E-F), except using disome footprints obtained from rich medium cultivated yeast cells.

(I) The aggregated abundance profile around the start codon (left) and stop codon (right) of the 5′-end of monosome footprints. The footprint abundance at each codon site was normalized by the total reads of the corresponding gene before aggregation.

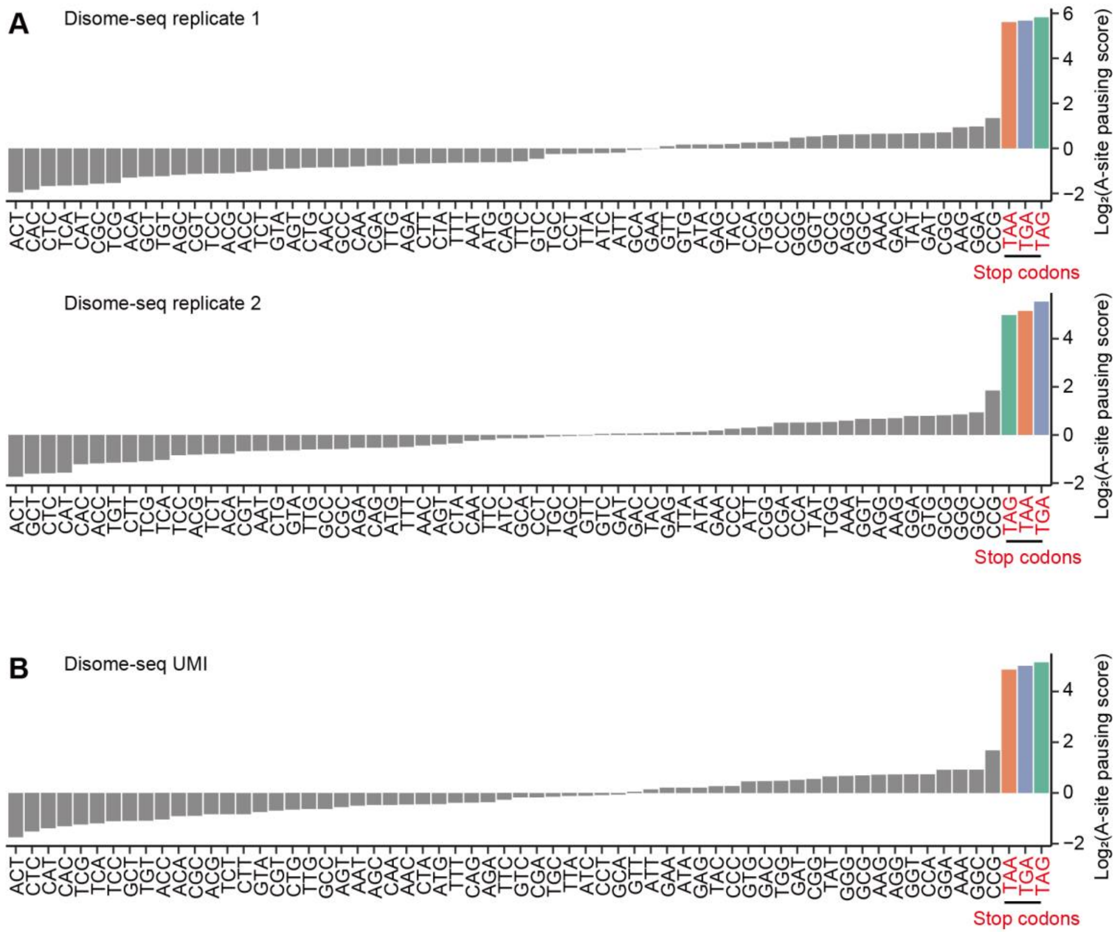


**Figure S4. The pausing signal at the A-site was consistent among different data processing approaches.**

(A) The A-site pausing scores for disome-footprint replicate 1 and disome-footprint replicate 2, respectively.

(B) The A-site pausing scores for disome footprints after removing PCR duplicates using UMI.

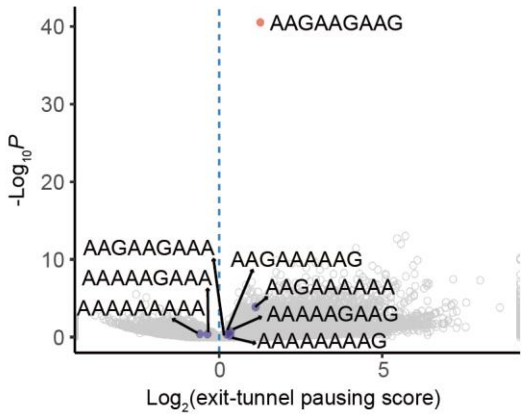


**Figure S5. Triple-lysine encoded by triple-AAG displayed the strongest collision signal**

Volcano plot shows the exit-tunnel pausing score for disome footprints of each of the $61^3$ codon 3-mers, and the corresponding *P*-value was given by the Mantel-Haenszel test. The codon 3-mers of triple-lysine are highlighted.

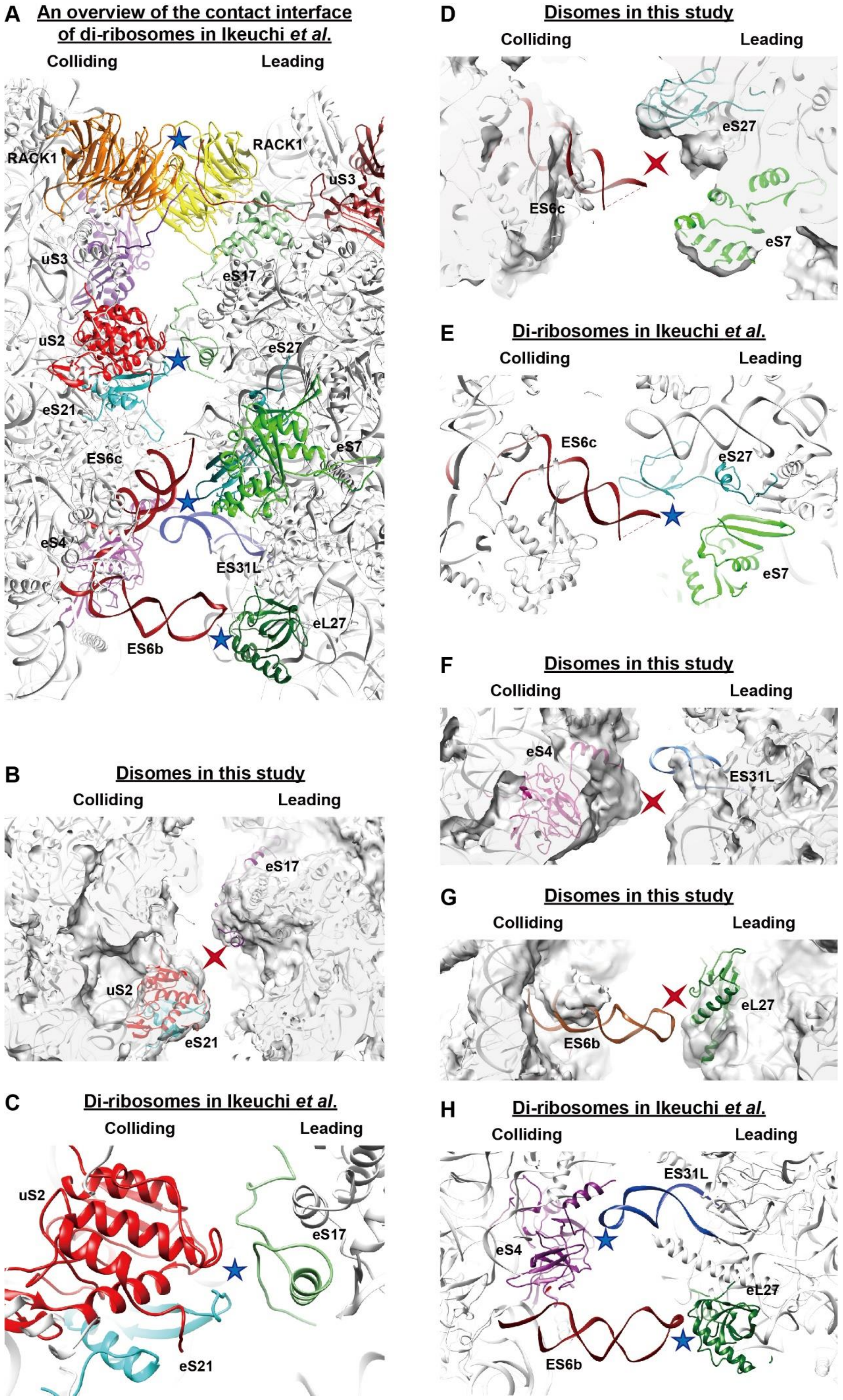
A An overview of the contact interface of di-ribosomes in Ikeuchi *et al.*
Colliding
Leading
RACK1
RACK1
uS3
uS3
eS17
uS2
eS27
eS21
ES6c
eS7
eS4
ES31L
eL27
ES6b
B Disomes in this study
Colliding
Leading
eS17
uS2
eS21
C Di-ribosomes in Ikeuchi *et al.*
Colliding
Leading
uS2
eS17
eS21
D Disomes in this study
Colliding
Leading
eS27
ES6c
eS7
E Di-ribosomes in Ikeuchi *et al.*
Colliding
Leading
ES6c
eS27
eS7
F Disomes in this study
Colliding
Leading
eS4
ES31L
G Disomes in this study
Colliding
Leading
eL27
ES6b
H Di-ribosomes in Ikeuchi *et al.*
Colliding
Leading
ES31L
eS4
eL27
ES6b

**Figure S6. The contact interface of disomes and the RQC-inducing di-ribosomes.**
(A) The zoomed-in overview of the contact interface between the 40S subunits of the RQC-inducing di-ribosomes (Ikeuchi *et al*. 2019).
(B-C) The zoomed-in view of the 40S body-to-body contact site of a disome (B) and a di-ribosome (C). The interactions between eS17 of the leading ribosome and the uS2 and eS21 of the colliding ribosome observed in di-ribosomes are absent in disomes.
(D-E) The zoomed-in view of the 40S platform-to-platform contact site of disomes (D) and di-ribosomes (E). The interactions between eS27, eS7 of the leading ribosome and the expansion segment ES6c of the colliding ribosome observed in di-ribosomes are absent in disomes.
(F-H) The zoomed-in view of the 60S-to-40S contact site of disomes (F-G) and di-ribosomes (H). The dual interactions between ES31L of the leading ribosome and eS4 in the colliding ribosome, together with eL27 and ES6c observed in di-ribosomes are absent in disomes. All blue stars indicate strong interactions between the two ribosomes in di-ribosomes. In contrast, these interactions are absent in disomes (marked by the red crosses).

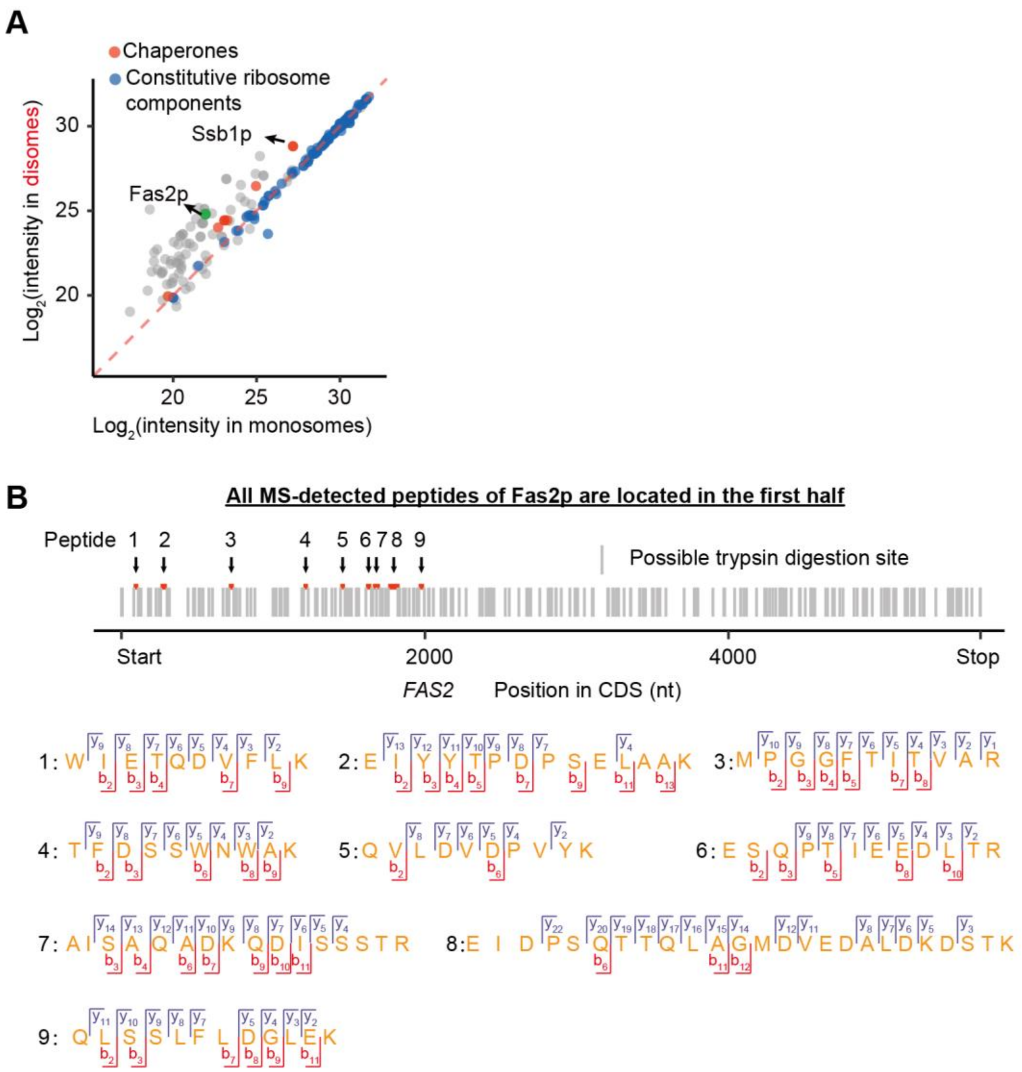


**Figure S7. The identification of disome-associated proteins.**

(A) The biological replicate of **Figure 6B**. Fas2p is highlighted in green.

(B) MS-detected peptides (red) and the N-termini of the theoretical peptides generated by *in silico* trypsin digestion (grey) of Fas2p. MS/MS spectra are shown.

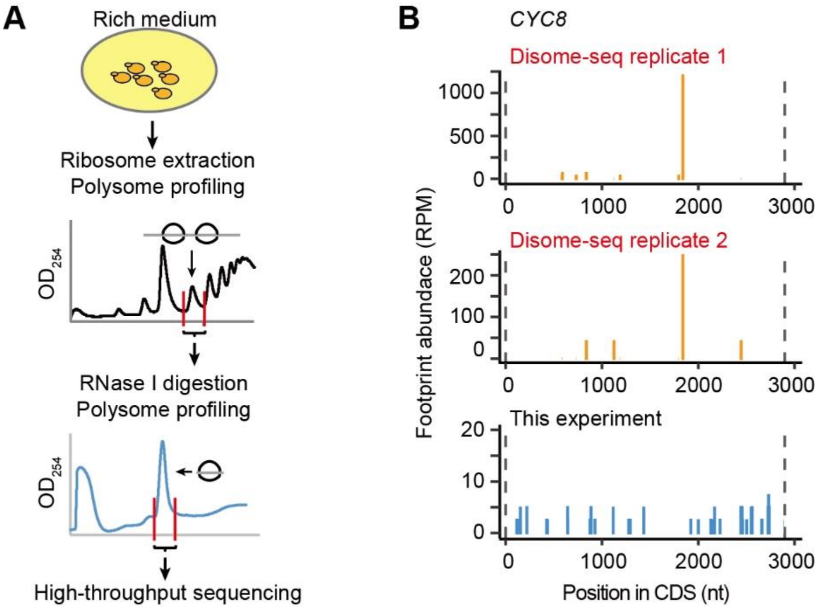


**Figure S8. Disomes were not from the two-ribosome-containing transcripts.**

(A) The ribosome footprints in the transcripts bound by two ribosomes were sequenced.

(B) The ribosome collision that was repeatedly detected in disome-seq (orange) was not observed in this experiment (blue).

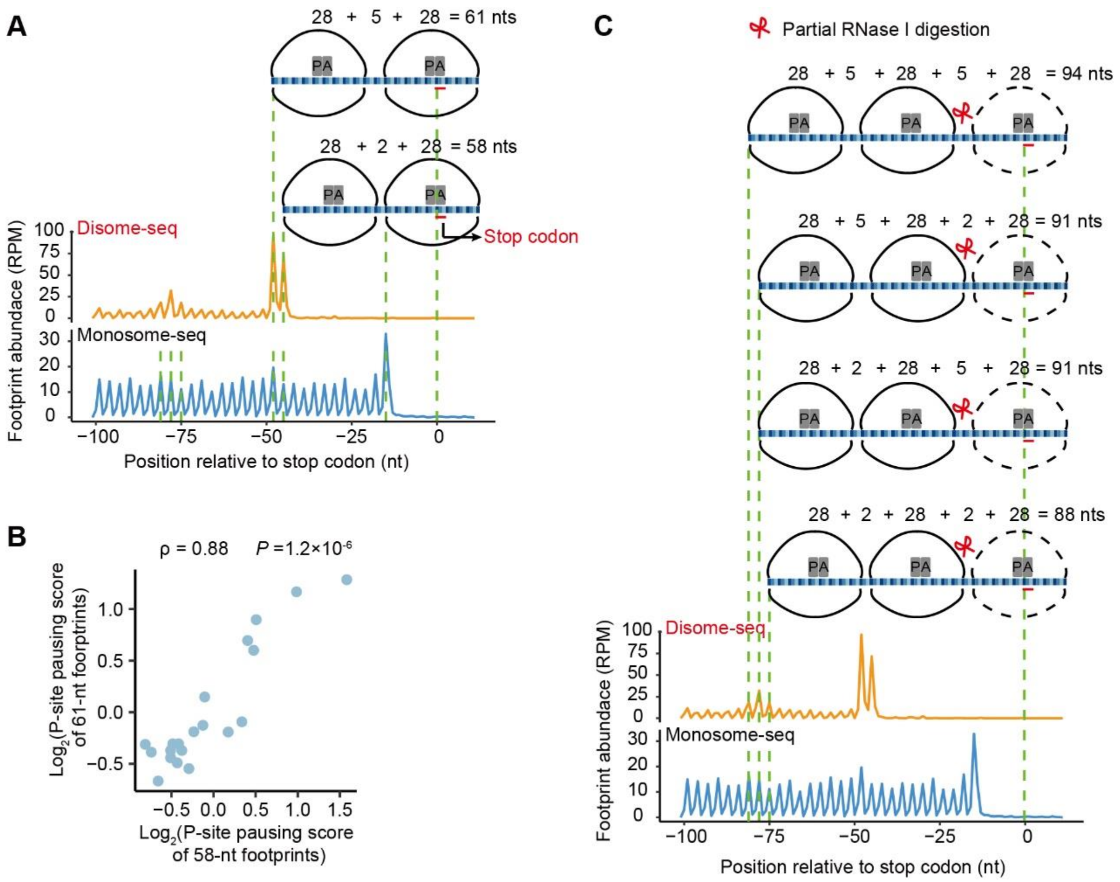


**Figure S9. The 58-nt and 61-nt disome footprints.**

(A) The aggregated abundance profile around the stop codon of the 5′-end of monosome (blue) and disome footprints (orange). The footprint abundance at each codon site was normalized by the total reads of the corresponding gene before aggregation. Two peaks were observed at the $45^{th}$ nt and $48^{th}$ nt upstream of the stop codon in the profile of disome footprints. The presumed conformations of the 58-nt disome and 61-nt disome are shown.

(B) The P-site pausing scores calculated from the 61-nt disome footprints are plotted against those from the 58-nt disome footprints. The *P*-value was given by Spearman's correlation.

(C) The conformations of trisomes, inferred from the footprints around the stop codon. Disome footprints with the 5′-end ~75-nt upstream of the stop codon were generated likely through the RNase digestion between the mid- and the 3′-ribosome in a trisome.

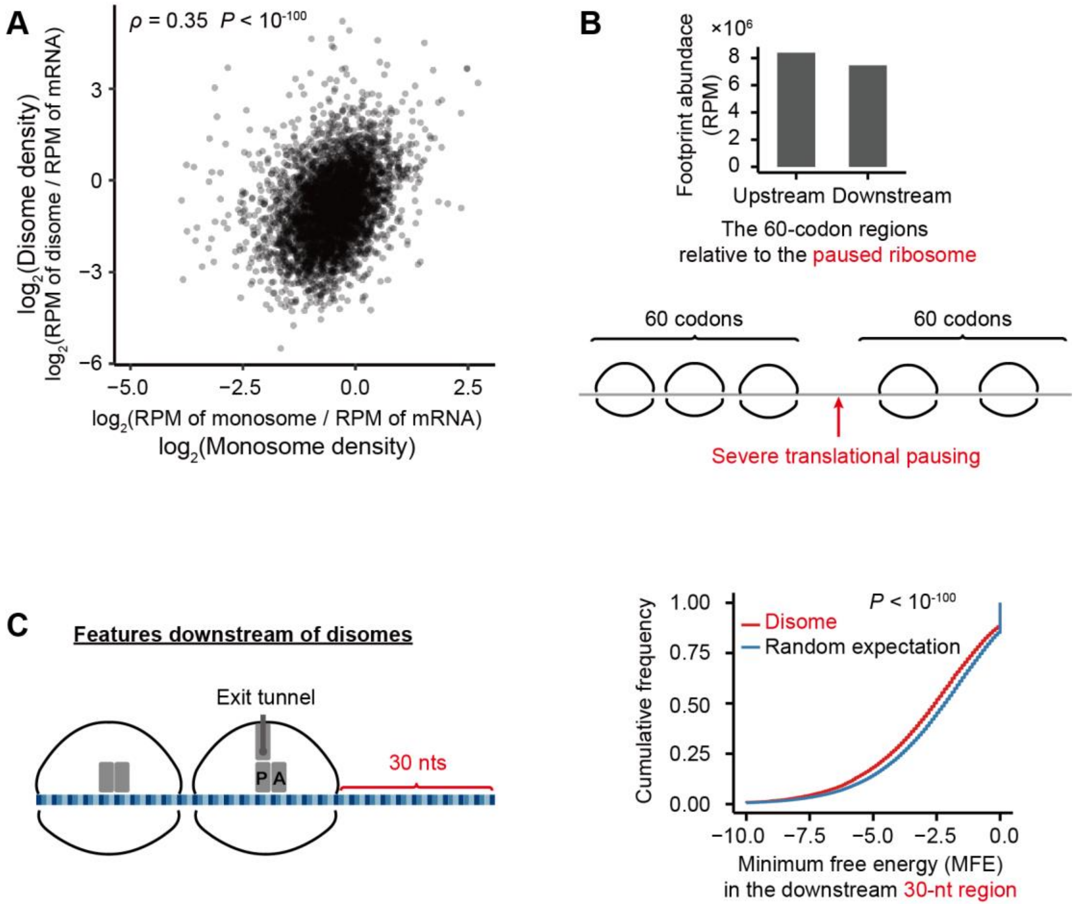


**Figure S10. Features upstream and downstream of the paused ribosomes.**

(A) Higher disome density was observed in the gene with higher monosome density.

(B) Lower ribosome density was observed downstream of the paused ribosomes.

(C) mRNA regions downstream of disomes exhibit stronger secondary structures than the random expectation. The red line shows the cumulative curve of the minimum free energy (MFE) of the 30-nt mRNA fragments downstream of disomes (red line). A random 30-nt mRNA fragment on the corresponding gene was used as a control (blue line). *P*-value was given by the Kolmogorov-Smirnov test.

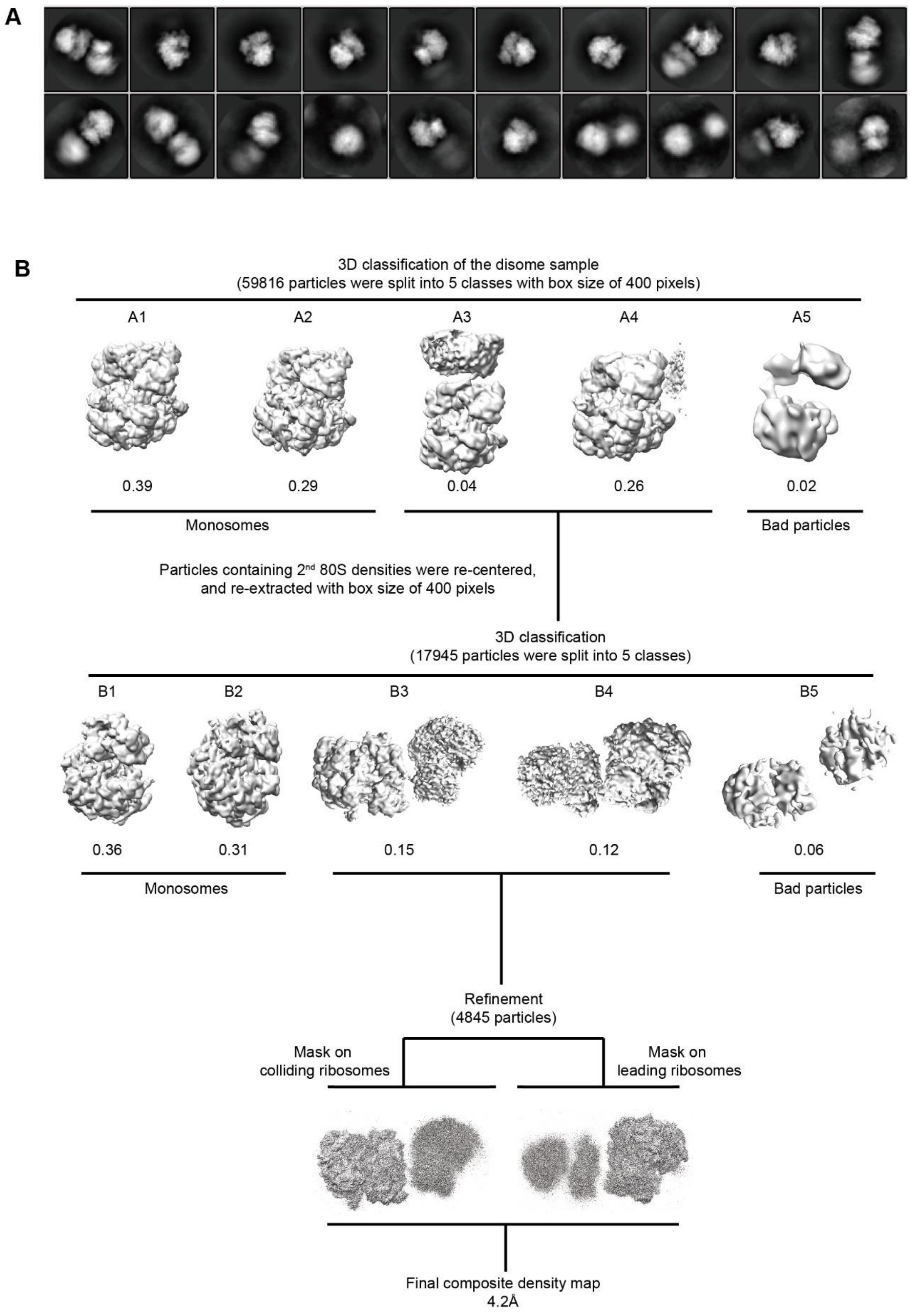


**Figure S11. Cryo-EM data processing of disome particles.**

(A) Representative 2D class averages of disome particles.

(B) The flow chart for 3D classification and refinement.

### Supplementary Tables

#### Table S1. Summary of the monosome-seq library.

| Library | Total reads | Filtered reads[a] | Uniquely mapped reads[b] |
|---|---|---|---|
| Replicate 1 | 24693292 | 2420449 | 420358 |
| Replicate 2 | 20833790 | 1659124 | 336258 |
| 3AT-replicate 1 | 15470359 | 1897389 | 157989 |
| 3AT-replicate 2 | 14574600 | 1675158 | 147562 |
| Control in Figure S8 | 26377014 | 5206211 | 858732 |

[a] Reads after removing rRNA.

[b] Reads uniquely mapped to the yeast genome.

#### Table S2. Summary of the disome-seq library.

| Library | Total reads | Filtered reads[a] | Uniquely mapped reads[b] | 50-75 bp reads |
|---|---|---|---|---|
| Replicate 1 | 272473666 | 13373574 | 2628459 | 100114 |
| Replicate 2 | 263337526 | 4410686 | 681055 | 55765 |
| 3AT- replicate 1 | 36139349 | 6144821 | 36084 | |
| 3AT- replicate 2 | 39727602 | 8068750 | 44211 | |

[a] Reads after removing rRNA.

[b] Reads uniquely mapped to the yeast genome.

#### Table S3. Summary of the mRNA-seq library.

| Library | Total reads | Uniquely mapped reads[a] |
|---|---|---|
| Replicate 1 | 5478547 | 1337661 |
| Replicate 2 | 6670236 | 1580266 |
| 3AT- replicate 1 | 6289058 | 1861188 |
| 3AT- replicate 2 | 4062390 | 1322674 |

[a] Reads uniquely mapped to the yeast genome.

#### Table S4. Disome-associated proteins (replicate 1: heavy isotope labeled disome proteins, disome/monosome intensity ratio > 1.5).

| ORF | Gene | Description* |
|---|---|---|
| **Chaperone** | | |
| YAL005C | *SSA1* | ATPase involved in protein folding and nuclear localization signal (NLS)-directed nuclear transport; member of heat shock protein 70 (HSP70) family; forms a chaperone complex with Ydj1p; localized to the nucleus, cytoplasm, and cell wall; 98% identical with Ssa2p, but subtle differences between the two proteins provide functional specificity with respect to propagation of yeast [URE3] prions and vacuolar-mediated degradations of gluconeogenesis enzymes |
| YLL024C | *SSA2* | ATP binding protein involved in protein folding and vacuolar import of proteins; member of heat shock protein 70 (HSP70) family; associated with the chaperonin-containing T-complex; present in the cytoplasm, vacuolar membrane and cell wall; 98% identical with |

| | | |
|---|---|---|
| | | Ssa1p, but subtle differences between the two proteins provide functional specificity with respect to propagation of yeast [URE3] prions and vacuolar-mediated degradations of gluconeogenesis enzymes |
| YDL229W | *SSB1* | Cytoplasmic ATPase that is a ribosome-associated molecular chaperone, functions with J-protein partner Zuo1p; may be involved in folding of newly-made polypeptide chains; member of the HSP70 family; interacts with phosphatase subunit Reg1p |
| YNL209W | *SSB2* | Cytoplasmic ATPase that is a ribosome-associated molecular chaperone, functions with J-protein partner Zuo1p; may be involved in the folding of newly-synthesized polypeptide chains; member of the HSP70 family; homolog of SSB1 |
| YMR186W | *HSC82* | Cytoplasmic chaperone of the Hsp90 family, redundant in function and nearly identical with Hsp82p, and together they are essential; expressed constitutively at 10-fold higher basal levels than HSP82 and induced 2-3 fold by heat shock |
| YPL240C | *HSP82* | Hsp90 chaperone required for pheromone signaling and negative regulation of Hsf1p; docks with Tom70p for mitochondrial preprotein delivery; promotes telomerase DNA binding and nucleotide addition; interacts with Cns1p, Cpr6p, Cpr7p, Sti1p |
| YJL159W | *HSP150* | O-mannosylated heat shock protein that is secreted and covalently attached to the cell wall via beta-1,3-glucan and disulfide bridges; required for cell wall stability; induced by heat shock, oxidative stress, and nitrogen limitation |
| YIL016W | *SNL1* | Protein of unknown function proposed to be involved in nuclear pore complex biogenesis and maintenance as well as protein folding; has similarity to the mammalian BAG-1 protein |
| YJL008C | *CCT8* | Subunit of the cytosolic chaperonin Cct ring complex, related to Tcp1p, required for the assembly of actin and tubulins in vivo |
| YDR033W | *MRH1* | Protein that localizes primarily to the plasma membrane, also found at the nuclear envelope; the authentic, non-tagged protein is detected in mitochondria in a phosphorylated state; has similarity to Hsp30p and Yro2p |
| YNL064C | *YDJ1* | Type I HSP40 co-chaperone involved in regulation of the HSP90 and HSP70 functions; involved in protein translocation across membranes; member of the DnaJ family |
| **Helicase** | | |
| YJL138C | *TIF2* | Translation initiation factor eIF4A, identical to Tif1p; DEA(D/H)-box RNA helicase that couples ATPase activity to RNA binding and unwinding; forms a dumbbell structure of two compact domains connected by a linker; interacts with eIF4G |
| YKR059W | *TIF1* | Translation initiation factor eIF4A, identical to Tif2p; DEA(D/H)-box RNA helicase that couples ATPase activity to RNA binding and |

| | | |
|---|---|---|
| | | unwinding; forms a dumbbell structure of two compact domains connected by a linker; interacts with eIF4G |
| **Metabolic** | | |
| YPL061W | *ALD6* | Cytosolic aldehyde dehydrogenase, activated by Mg2+ and utilizes NADP+ as the preferred coenzyme; required for conversion of acetaldehyde to acetate; constitutively expressed; locates to the mitochondrial outer surface upon oxidative stress |
| YGR192C | *TDH3* | Glyceraldehyde-3-phosphate dehydrogenase, isozyme 3, involved in glycolysis and gluconeogenesis; tetramer that catalyzes the reaction of glyceraldehyde-3-phosphate to 1,3 bis-phosphoglycerate; detected in the cytoplasm and cell wall |
| YJR009C | *TDH2* | Glyceraldehyde-3-phosphate dehydrogenase, isozyme 2, involved in glycolysis and gluconeogenesis; tetramer that catalyzes the reaction of glyceraldehyde-3-phosphate to 1,3 bis-phosphoglycerate; detected in the cytoplasm and cell wall |
| YOL086C | *ADH1* | Alcohol dehydrogenase, fermentative isozyme active as homo- or heterotetramers; required for the reduction of acetaldehyde to ethanol, the last step in the glycolytic pathway |
| YDR502C | *SAM2* | S-adenosylmethionine synthetase, catalyzes transfer of the adenosyl group of ATP to the sulfur atom of methionine; one of two differentially regulated isozymes (Sam1p and Sam2p) |
| YLR180W | *SAM1* | S-adenosylmethionine synthetase, catalyzes transfer of the adenosyl group of ATP to the sulfur atom of methionine; one of two differentially regulated isozymes (Sam1p and Sam2p) |
| YLR044C | *PDC1* | Major of three pyruvate decarboxylase isozymes, key enzyme in alcoholic fermentation, decarboxylates pyruvate to acetaldehyde; subject to glucose-, ethanol-, and autoregulation; involved in amino acid catabolism |
| YPL231W | *FAS2* | Alpha subunit of fatty acid synthetase, which catalyzes the synthesis of long-chain saturated fatty acids; contains the acyl-carrier protein domain and beta-ketoacyl reductase, beta-ketoacyl synthase and self-pantetheinylation activities |
| YJL130C | *URA2* | Bifunctional carbamoylphosphate synthetase (CPSase)-aspartate transcarbamylase (ATCase), catalyzes the first two enzymatic steps in the de novo biosynthesis of pyrimidines; both activities are subject to feedback inhibition by UTP |
| YBR196C | *PGI1* | Glycolytic enzyme phosphoglucose isomerase, catalyzes the interconversion of glucose-6-phosphate and fructose-6-phosphate; required for cell cycle progression and completion of the gluconeogenic events of sporulation |
| YKL060C | *FBA1* | Fructose 1,6-bisphosphate aldolase, required for glycolysis and gluconeogenesis; catalyzes conversion of fructose 1,6 bisphosphate to glyceraldehyde-3-P and dihydroxyacetone-P; locates to mitochondrial outer surface upon oxidative stress |

| | | |
|---|---|---|
| YAL038W | *CDC19* | Pyruvate kinase, functions as a homotetramer in glycolysis to convert phosphoenolpyruvate to pyruvate, the input for aerobic (TCA cycle) or anaerobic (glucose fermentation) respiration |
| YKL182W | *FAS1* | Beta subunit of fatty acid synthetase, which catalyzes the synthesis of long-chain saturated fatty acids; contains acetyltransacylase, dehydratase, enoyl reductase, malonyl transacylase, and palmitoyl transacylase activities |
| YKL152C | *GPM1* | Tetrameric phosphoglycerate mutase, mediates the conversion of 3-phosphoglycerate to 2-phosphoglycerate during glycolysis and the reverse reaction during gluconeogenesis |
| YIL078W | *THS1* | Threonyl-tRNA synthetase, essential cytoplasmic protein |
| YGR254W | *ENO1* | Enolase I, a phosphopyruvate hydratase that catalyzes the conversion of 2-phosphoglycerate to phosphoenolpyruvate during glycolysis and the reverse reaction during gluconeogenesis; expression is repressed in response to glucose |
| YHR174W | *ENO2* | Enolase II, a phosphopyruvate hydratase that catalyzes the conversion of 2-phosphoglycerate to phosphoenolpyruvate during glycolysis and the reverse reaction during gluconeogenesis; expression is induced in response to glucose |
| YMR205C | *PFK2* | Beta subunit of heterooctameric phosphofructokinase involved in glycolysis, indispensable for anaerobic growth, activated by fructose-2,6-bisphosphate and AMP, mutation inhibits glucose induction of cell cycle-related genes |
| YHR019C | *DED81* | Cytosolic asparaginyl-tRNA synthetase, required for protein synthesis, catalyzes the specific attachment of asparagine to its cognate tRNA |
| YJL153C | *INO1* | Inositol-3-phosphate synthase, involved in synthesis of inositol phosphates and inositol-containing phospholipids; transcription is coregulated with other phospholipid biosynthetic genes by Ino2p and Ino4p, which bind the UASINO DNA element |
| YLR354C | *TAL1* | Transaldolase, enzyme in the non-oxidative pentose phosphate pathway; converts sedoheptulose 7-phosphate and glyceraldehyde 3-phosphate to erythrose 4-phosphate and fructose 6-phosphate |
| YDR050C | *TPI1* | Triose phosphate isomerase, abundant glycolytic enzyme; mRNA half-life is regulated by iron availability; transcription is controlled by activators Reb1p, Gcr1p, and Rap1p through binding sites in the 5′ non-coding region |
| YNR016C | *ACC1* | Acetyl-CoA carboxylase, biotin containing enzyme that catalyzes the carboxylation of acetyl-CoA to form malonyl-CoA; required for de novo biosynthesis of long-chain fatty acids |
| YCR012W | *PGK1* | 3-phosphoglycerate kinase, catalyzes transfer of high-energy phosphoryl groups from the acyl phosphate of 1,3-bisphosphoglycerate to ADP to produce ATP; key enzyme in glycolysis and gluconeogenesis |

| **Other** | | |
|---|---|---|
| YMR307W | *GAS1* | Beta-1,3-glucanosyltransferase, required for cell wall assembly and also has a role in transcriptional silencing; localizes to the cell surface via a glycosylphosphatidylinositol (GPI) anchor; also found at the nuclear periphery |
| YGR282C | *BGL2* | Endo-beta-1,3-glucanase, major protein of the cell wall, involved in cell wall maintenance |
| YFL039C | *ACT1* | Actin, structural protein involved in cell polarization, endocytosis, and other cytoskeletal functions |
| YDL055C | *PSA1* | GDP-mannose pyrophosphorylase (mannose-1-phosphate guanyltransferase), synthesizes GDP-mannose from GTP and mannose-1-phosphate in cell wall biosynthesis; required for normal cell wall structure |
| YKL164C | *PIR1* | O-glycosylated protein required for cell wall stability; attached to the cell wall via beta-1,3-glucan; mediates mitochondrial translocation of Apn1p; expression regulated by the cell integrity pathway and by Swi5p during the cell cycle |
| YBR009C | *HHF1* | Histone H4, core histone protein required for chromatin assembly and chromosome function; one of two identical histone proteins (see also HHF2); contributes to telomeric silencing; N-terminal domain involved in maintaining genomic integrity |
| YNL030W | *HHF2* | Histone H4, core histone protein required for chromatin assembly and chromosome function; one of two identical histone proteins (see also HHF1); contributes to telomeric silencing; N-terminal domain involved in maintaining genomic integrity |
| YBL002W | *HTB2* | Histone H2B, core histone protein required for chromatin assembly and chromosome function; nearly identical to HTB1; Rad6p-Bre1p-Lge1p mediated ubiquitination regulates transcriptional activation, meiotic DSB formation and H3 methylation |
| YDR224C | *HTB1* | Histone H2B, core histone protein required for chromatin assembly and chromosome function; nearly identical to HTB2; Rad6p-Bre1p-Lge1p mediated ubiquitination regulates transcriptional activation, meiotic DSB formation and H3 methylation |
| YFL037W | *TUB2* | Beta-tubulin; associates with alpha-tubulin (Tub1p and Tub3p) to form tubulin dimer, which polymerizes to form microtubules |
| YNL284C | *MRPL10* | Mitochondrial ribosomal protein of the large subunit; appears as two protein spots (YmL10 and YmL18) on two-dimensional SDS gels |
| YDL202W | *MRPL11* | Mitochondrial ribosomal protein of the large subunit |
| YPL093W | *NOG1* | Putative GTPase that associates with free 60S ribosomal subunits in the nucleolus and is required for 60S ribosomal subunit biogenesis; constituent of 66S pre-ribosomal particles; member of the ODN family of nucleolar G-proteins |

| | | |
|---|---|---|
| YPR016C | *TIF6* | Constituent of 66S pre-ribosomal particles, has similarity to human translation initiation factor 6 (eIF6); may be involved in the biogenesis and or stability of 60S ribosomal subunits |
| YER006W | *NUG1* | GTPase that associates with nuclear 60S pre-ribosomes, required for export of 60S ribosomal subunits from the nucleus |
| YHR088W | *RPF1* | Nucleolar protein involved in the assembly and export of the large ribosomal subunit; constituent of 66S pre-ribosomal particles; contains a sigma(70)-like motif, which is thought to bind RNA |
| YMR012W | *CLU1* | eIF3 component of unknown function; deletion causes defects in mitochondrial organization but not in growth or translation initiation, can rescue cytokinesis and mitochondrial organization defects of the Dictyostelium cluA- mutant |
| YDR385W | *EFT2* | Elongation factor 2 (EF-2), also encoded by EFT1; catalyzes ribosomal translocation during protein synthesis; contains diphthamide, the unique posttranslationally modified histidine residue specifically ADP-ribosylated by diphtheria toxin |
| YOR133W | *EFT1* | Elongation factor 2 (EF-2), also encoded by EFT2; catalyzes ribosomal translocation during protein synthesis; contains diphthamide, the unique posttranslationally modified histidine residue specifically ADP-ribosylated by diphtheria toxin |
| YLR249W | *YEF3* | Gamma subunit of translational elongation factor eEF1B, stimulates the binding of aminoacyl-tRNA (AA-tRNA) to ribosomes by releasing eEF1A (Tef1p/Tef2p) from the ribosomal complex; contains two ABC cassettes; binds and hydrolyzes ATP |
| YBR118W | *TEF2* | Translational elongation factor EF-1 alpha; also encoded by TEF1; functions in the binding reaction of aminoacyl-tRNA (AA-tRNA) to ribosomes; may also have a role in tRNA re-export from the nucleus |
| YPR080W | *TEF1* | Translational elongation factor EF-1 alpha; also encoded by TEF2; functions in the binding reaction of aminoacyl-tRNA (AA-tRNA) to ribosomes; may also have a role in tRNA re-export from the nucleus |
| YGL008C | *PMA1* | Plasma membrane H+-ATPase, pumps protons out of the cell; major regulator of cytoplasmic pH and plasma membrane potential; P2-type ATPase; Hsp30p plays a role in Pma1p regulation; interactions with Std1p appear to propagate [GAR+] |
| YPL036W | *PMA2* | Plasma membrane H+-ATPase, isoform of Pma1p, involved in pumping protons out of the cell; regulator of cytoplasmic pH and plasma membrane potential |
| YBR127C | *VMA2* | Subunit B of the eight-subunit V1 peripheral membrane domain of the vacuolar H+-ATPase (V-ATPase), an electrogenic proton pump found throughout the endomembrane system; contains nucleotide binding sites; also detected in the cytoplasm |
| YDL185W | *VMA1* | Subunit A of the eight-subunit V1 peripheral membrane domain of the vacuolar H+-ATPase; protein precursor undergoes self-catalyzed |

| | | |
|---|---|---|
| | | splicing to yield the extein Tfp1p and the intein Vde (PI-SceI), which is a site-specific endonuclease |
| YDR233C | *RTN1* | ER membrane protein that interacts with Sey1p to maintain ER morphology; interacts with exocyst subunit Sec6p, with Yip3p, and with Sbh1p; null mutant has an altered ER morphology; member of the RTNLA (reticulon-like A) subfamily |

* Descriptions were retrieved from the Saccharomyces Genome Database (https://www.yeastgenome.org/)

**Table S5. Disome-associated proteins (replicate 2: light isotope labeled disome proteins, disome/monosome intensity ratio > 1.5).**

| ORF | Gene | Description* |
|---|---|---|
| **Chaperone** | | |
| YAL005C | *SSA1* | ATPase involved in protein folding and nuclear localization signal (NLS)-directed nuclear transport; member of heat shock protein 70 (HSP70) family; forms a chaperone complex with Ydj1p; localized to the nucleus, cytoplasm, and cell wall; 98% identical with Ssa2p, but subtle differences between the two proteins provide functional specificity with respect to propagation of yeast [URE3] prions and vacuolar-mediated degradations of gluconeogenesis enzymes |
| YLL024C | *SSA2* | ATP binding protein involved in protein folding and vacuolar import of proteins; member of heat shock protein 70 (HSP70) family; associated with the chaperonin-containing T-complex; present in the cytoplasm, vacuolar membrane and cell wall; 98% identical with Ssa1p, but subtle differences between the two proteins provide functional specificity with respect to propagation of yeast [URE3] prions and vacuolar-mediated degradations of gluconeogenesis enzymes |
| YDL229W | *SSB1* | Cytoplasmic ATPase that is a ribosome-associated molecular chaperone, functions with J-protein partner Zuo1p; may be involved in folding of newly-made polypeptide chains; member of the HSP70 family; interacts with phosphatase subunit Reg1p |
| YNL209W | *SSB2* | Cytoplasmic ATPase that is a ribosome-associated molecular chaperone, functions with J-protein partner Zuo1p; may be involved in the folding of newly-synthesized polypeptide chains; member of the HSP70 family; homolog of SSB1 |
| YMR186W | *HSC82* | Cytoplasmic chaperone of the Hsp90 family, redundant in function and nearly identical with Hsp82p, and together they are essential; expressed constitutively at 10-fold higher basal levels than HSP82 and induced 2-3 fold by heat shock |
| YPL240C | *HSP82* | Hsp90 chaperone required for pheromone signaling and negative regulation of Hsf1p; docks with Tom70p for mitochondrial preprotein delivery; promotes telomerase DNA binding and nucleotide addition; interacts with Cns1p, Cpr6p, Cpr7p, Sti1p |
| YJL159W | *HSP150* | O-mannosylated heat shock protein that is secreted and covalently attached to the cell wall via beta-1,3-glucan and disulfide bridges; required for cell wall stability; induced by heat shock, oxidative stress, and nitrogen limitation |
| **Helicase** | | |
| YJL138C | *TIF2* | Translation initiation factor eIF4A, identical to Tif1p; DEA(D/H)-box RNA helicase that couples ATPase activity to RNA binding and unwinding; forms a dumbbell structure of two compact domains connected by a linker; interacts with eIF4G |

| | | |
|---|---|---|
| YKR059W | *TIF1* | Translation initiation factor eIF4A, identical to Tif2p; DEA(D/H)-box RNA helicase that couples ATPase activity to RNA binding and unwinding; forms a dumbbell structure of two compact domains connected by a linker; interacts with eIF4G |
| YLR419W | YLR419W | Putative helicase with limited sequence similarity to human Rb protein; the authentic, non-tagged protein is detected in highly purified mitochondria in high-throughput studies; YLR419W is not an essential gene |
| **Metabolic** | | |
| YOL086C | *ADH1* | Alcohol dehydrogenase, fermentative isozyme active as homo- or heterotetramers; required for the reduction of acetaldehyde to ethanol, the last step in the glycolytic pathway |
| YJL130C | *URA2* | Bifunctional carbamoylphosphate synthetase (CPSase)-aspartate transcarbamylase (ATCase), catalyzes the first two enzymatic steps in the de novo biosynthesis of pyrimidines; both activities are subject to feedback inhibition by UTP |
| YJR009C | *TDH2* | Glyceraldehyde-3-phosphate dehydrogenase, isozyme 2, involved in glycolysis and gluconeogenesis; tetramer that catalyzes the reaction of glyceraldehyde-3-phosphate to 1,3 bis-phosphoglycerate; detected in the cytoplasm and cell wall |
| YDR502C | *SAM2* | S-adenosylmethionine synthetase, catalyzes transfer of the adenosyl group of ATP to the sulfur atom of methionine; one of two differentially regulated isozymes (Sam1p and Sam2p) |
| YLR180W | *SAM1* | S-adenosylmethionine synthetase, catalyzes transfer of the adenosyl group of ATP to the sulfur atom of methionine; one of two differentially regulated isozymes (Sam1p and Sam2p) |
| YCL064C | *CHA1* | Catabolic L-serine (L-threonine) deaminase, catalyzes the degradation of both L-serine and L-threonine; required to use serine or threonine as the sole nitrogen source, transcriptionally induced by serine and threonine |
| YKL060C | *FBA1* | Fructose 1,6-bisphosphate aldolase, required for glycolysis and gluconeogenesis; catalyzes conversion of fructose 1,6 bisphosphate to glyceraldehyde-3-P and dihydroxyacetone-P; locates to mitochondrial outer surface upon oxidative stress |
| YLR044C | *PDC1* | Major of three pyruvate decarboxylase isozymes, key enzyme in alcoholic fermentation, decarboxylates pyruvate to acetaldehyde; subject to glucose-, ethanol-, and autoregulation; involved in amino acid catabolism |
| YKL182W | *FAS1* | Beta subunit of fatty acid synthetase, which catalyzes the synthesis of long-chain saturated fatty acids; contains acetyltransacylase, dehydratase, enoyl reductase, malonyl transacylase, and palmitoyl transacylase activities |
| YBR196C | *PGI1* | Glycolytic enzyme phosphoglucose isomerase, catalyzes the interconversion of glucose-6-phosphate and fructose-6-phosphate; |

| | | |
|---|---|---|
| | | required for cell cycle progression and completion of the gluconeogenic events of sporulation |
| YJR073C | *OPI3* | Phospholipid methyltransferase (methylene-fatty-acyl-phospholipid synthase), catalyzes the last two steps in phosphatidylcholine biosynthesis |
| YIL078W | *THS1* | Threonyl-tRNA synthetase, essential cytoplasmic protein |
| YPL231W | *FAS2* | Alpha subunit of fatty acid synthetase, which catalyzes the synthesis of long-chain saturated fatty acids; contains the acyl-carrier protein domain and beta-ketoacyl reductase, beta-ketoacyl synthase and self-pantetheinylation activities |
| YER091C | *MET6* | Cobalamin-independent methionine synthase, involved in methionine biosynthesis and regeneration; requires a minimum of two glutamates on the methyltetrahydrofolate substrate, similar to bacterial metE homologs |
| YAL038W | *CDC19* | Pyruvate kinase, functions as a homotetramer in glycolysis to convert phosphoenolpyruvate to pyruvate, the input for aerobic (TCA cycle) or anaerobic (glucose fermentation) respiration |
| YHR174W | *ENO2* | Enolase II, a phosphopyruvate hydratase that catalyzes the conversion of 2-phosphoglycerate to phosphoenolpyruvate during glycolysis and the reverse reaction during gluconeogenesis; expression is induced in response to glucose |
| YLR355C | *ILV5* | Bifunctional acetohydroxyacid reductoisomerase and mtDNA binding protein; involved in branched-chain amino acid biosynthesis and maintenance of wild-type mitochondrial DNA; found in mitochondrial nucleoids |
| YMR205C | *PFK2* | Beta subunit of heterooctameric phosphofructokinase involved in glycolysis, indispensable for anaerobic growth, activated by fructose-2,6-bisphosphate and AMP, mutation inhibits glucose induction of cell cycle-related genes |
| YKL152C | *GPM1* | Tetrameric phosphoglycerate mutase, mediates the conversion of 3-phosphoglycerate to 2-phosphoglycerate during glycolysis and the reverse reaction during gluconeogenesis |
| YPR184W | *GDB1* | Glycogen debranching enzyme containing glucanotranferase and alpha-1,6-amyloglucosidase activities, required for glycogen degradation; phosphorylated in mitochondria; activity is inhibited by Igd1p |
| YCR012W | *PGK1* | 3-phosphoglycerate kinase, catalyzes transfer of high-energy phosphoryl groups from the acyl phosphate of 1,3-bisphosphoglycerate to ADP to produce ATP; key enzyme in glycolysis and gluconeogenesis |
| **Other** | | |
| YDL055C | *PSA1* | GDP-mannose pyrophosphorylase (mannose-1-phosphate guanyltransferase), synthesizes GDP-mannose from GTP and |

| | | |
|---|---|---|
| | | mannose-1-phosphate in cell wall biosynthesis; required for normal cell wall structure |
| YBR078W | *ECM33* | GPI-anchored protein of unknown function, has a possible role in apical bud growth; GPI-anchoring on the plasma membrane crucial to function; phosphorylated in mitochondria; similar to Sps2p and Pst1p |
| YGR282C | *BGL2* | Endo-beta-1,3-glucanase, major protein of the cell wall, involved in cell wall maintenance |
| YGR279C | *SCW4* | Cell wall protein with similarity to glucanases; scw4 scw10 double mutants exhibit defects in mating |
| YKL164C | *PIR1* | O-glycosylated protein required for cell wall stability; attached to the cell wall via beta-1,3-glucan; mediates mitochondrial translocation of Apn1p; expression regulated by the cell integrity pathway and by Swi5p during the cell cycle |
| YBR009C | *HHF1* | Histone H4, core histone protein required for chromatin assembly and chromosome function; one of two identical histone proteins (see also HHF2); contributes to telomeric silencing; N-terminal domain involved in maintaining genomic integrity |
| YNL030W | *HHF2* | Histone H4, core histone protein required for chromatin assembly and chromosome function; one of two identical histone proteins (see also HHF1); contributes to telomeric silencing; N-terminal domain involved in maintaining genomic integrity |
| YFL037W | *TUB2* | Beta-tubulin; associates with alpha-tubulin (Tub1p and Tub3p) to form tubulin dimer, which polymerizes to form microtubules |
| YFL039C | *ACT1* | Actin, structural protein involved in cell polarization, endocytosis, and other cytoskeletal functions |
| YJR113C | *RSM7* | Mitochondrial ribosomal protein of the small subunit, has similarity to E. coli S7 ribosomal protein |
| YPL093W | *NOG1* | Putative GTPase that associates with free 60S ribosomal subunits in the nucleolus and is required for 60S ribosomal subunit biogenesis; constituent of 66S pre-ribosomal particles; member of the ODN family of nucleolar G-proteins |
| YER006W | *NUG1* | GTPase that associates with nuclear 60S pre-ribosomes, required for export of 60S ribosomal subunits from the nucleus |
| YPR016C | *TIF6* | Constituent of 66S pre-ribosomal particles, has similarity to human translation initiation factor 6 (eIF6); may be involved in the biogenesis and or stability of 60S ribosomal subunits |
| YHR088W | *RPF1* | Nucleolar protein involved in the assembly and export of the large ribosomal subunit; constituent of 66S pre-ribosomal particles; contains a sigma(70)-like motif, which is thought to bind RNA |
| YAL025C | *MAK16* | Essential nuclear protein, constituent of 66S pre-ribosomal particles; required for maturation of 25S and 5.8S rRNAs; required for maintenance of M1 satellite double-stranded RNA of the L-A virus |

| | | |
|---|---|---|
| YGR103W | *NOP7* | Component of several different pre-ribosomal particles; forms a complex with Ytm1p and Erb1p that is required for maturation of the large ribosomal subunit; required for exit from G<sub0</sub and the initiation of cell proliferation |
| YHR072W-A | *NOP10* | Constituent of small nucleolar ribonucleoprotein particles containing H/ACA-type snoRNAs, which are required for pseudouridylation and processing of pre-18S rRNA |
| YOL077C | *BRX1* | Nucleolar protein, constituent of 66S pre-ribosomal particles; depletion leads to defects in rRNA processing and a block in the assembly of large ribosomal subunits; possesses a sigma(70)-like RNA-binding motif |
| YLR196W | *PWP1* | Protein with WD-40 repeats involved in rRNA processing; associates with trans-acting ribosome biogenesis factors; similar to beta-transducin superfamily |
| YLR175W | *CBF5* | Pseudouridine synthase catalytic subunit of box H/ACA small nucleolar ribonucleoprotein particles (snoRNPs), acts on both large and small rRNAs and on snRNA U2; mutations in human ortholog dyskerin cause the disorder dyskeratosis congenita |
| YPL061W | *ALD6* | Cytosolic aldehyde dehydrogenase, activated by Mg2+ and utilizes NADP+ as the preferred coenzyme; required for conversion of acetaldehyde to acetate; constitutively expressed; locates to the mitochondrial outer surface upon oxidative stress |
| YLR249W | *YEF3* | Gamma subunit of translational elongation factor eEF1B, stimulates the binding of aminoacyl-tRNA (AA-tRNA) to ribosomes by releasing eEF1A (Tef1p/Tef2p) from the ribosomal complex; contains two ABC cassettes; binds and hydrolyzes ATP |
| YKL081W | *TEF4* | Gamma subunit of translational elongation factor eEF1B, stimulates the binding of aminoacyl-tRNA (AA-tRNA) to ribosomes by releasing eEF1A (Tef1p/Tef2p) from the ribosomal complex |
| YDR385W | *EFT2* | Elongation factor 2 (EF-2), also encoded by EFT1; catalyzes ribosomal translocation during protein synthesis; contains diphthamide, the unique posttranslationally modified histidine residue specifically ADP-ribosylated by diphtheria toxin |
| YOR133W | *EFT1* | Elongation factor 2 (EF-2), also encoded by EFT2; catalyzes ribosomal translocation during protein synthesis; contains diphthamide, the unique posttranslationally modified histidine residue specifically ADP-ribosylated by diphtheria toxin |
| YMR012W | *CLU1* | eIF3 component of unknown function; deletion causes defects in mitochondrial organization but not in growth or translation initiation, can rescue cytokinesis and mitochondrial organization defects of the Dictyostelium cluA- mutant |
| YBR118W | *TEF2* | Translational elongation factor EF-1 alpha; also encoded by TEF1; functions in the binding reaction of aminoacyl-tRNA (AA-tRNA) to |

| | | |
|---|---|---|
| | | ribosomes; may also have a role in tRNA re-export from the nucleus |
| YPR080W | *TEF1* | Translational elongation factor EF-1 alpha; also encoded by TEF2; functions in the binding reaction of aminoacyl-tRNA (AA-tRNA) to ribosomes; may also have a role in tRNA re-export from the nucleus |
| YMR307W | *GAS1* | Beta-1,3-glucanosyltransferase, required for cell wall assembly and also has a role in transcriptional silencing; localizes to the cell surface via a glycosylphosphatidylinositol (GPI) anchor; also found at the nuclear periphery |
| YGL008C | *PMA1* | Plasma membrane H+-ATPase, pumps protons out of the cell; major regulator of cytoplasmic pH and plasma membrane potential; P2-type ATPase; Hsp30p plays a role in Pma1p regulation; interactions with Std1p appear to propagate [GAR+] |
| YPL036W | *PMA2* | Plasma membrane H+-ATPase, isoform of Pma1p, involved in pumping protons out of the cell; regulator of cytoplasmic pH and plasma membrane potential |
| YJR121W | *ATP2* | Beta subunit of the F1 sector of mitochondrial F1F0 ATP synthase, which is a large, evolutionarily conserved enzyme complex required for ATP synthesis; phosphorylated |
| YDL185W | *VMA1* | Subunit A of the eight-subunit V1 peripheral membrane domain of the vacuolar H+-ATPase; protein precursor undergoes self-catalyzed splicing to yield the extein Tfp1p and the intein Vde (PI-SceI), which is a site-specific endonuclease |

* Descriptions were retrieved from the Saccharomyces Genome Database (https://www.yeastgenome.org/)
